# Nested TAD hierarchy defines cohesion zones on replicated chromosomes

**DOI:** 10.64898/2026.05.02.722390

**Authors:** Zsuzsanna Takacs, Dmitry Mylarshchikov, Sofia Kolesnikova, Christoph C. H. Langer, Anton Goloborodko, Daniel W. Gerlich

## Abstract

During each cell cycle, cells must not only duplicate DNA sequence but also preserve the three-dimensional genome architecture required for gene regulation and DNA repair. After replication, this requires coordination between cohesin-mediated loop extrusion and sister-chromatid cohesion for homology-directed DNA repair. Because extrusion promotes sister resolution whereas cohesion can impede extrusion, how both activities coexist on replicated chromosomes is unclear. Here we show that cohesin function is partitioned within the nested TAD hierarchy: cohesive cohesin accumulates at high-level boundaries to form “cohesion zones,” while loop-extruding cohesin occupies boundaries across all hierarchical levels. Our data support a model in which this segregation emerges from dynamic interplay between cohesin pools and semi-permeable CTCF barriers, which act in cis and in trans to constrain sister–sister misalignment. Thus, a CTCF-defined boundary framework enables sister tethering while preserving dynamic loop folding, allowing replicated genomes to simultaneously support gene regulation and faithful DNA repair.

## INTRODUCTION

During each cell cycle, proliferating cells duplicate their genomes to pass an identical copy to daughter cells. This requires more than copying DNA sequence, because replicated sister chromatids must also preserve or re-establish three-dimensional folding and regulatory contact patterns that support genome function. In G2 phase, sister chromatids remain physically linked for homology-directed DNA repair and faithful chromosome segregation in the subsequent mitosis, while chromatin continues to form dynamic loops that support long-range regulation^1,2^. How replicated chromosomes coordinate persistent sister linkage with dynamic 3D folding, however, remains poorly understood.

A central mediator of 3D chromosome organization is cohesin^3,4^, a structural maintenance of chromosomes (SMC) complex with two mechanistically distinct activities. Through ATPase-dependent loop extrusion^5,6^, cohesin dynamically folds chromatin into loops and topologically associating domains (TADs)^7–13^, thereby regulating long - range enhancer – promoter communication^14–16^. Following DNA replication, a subset of cohesin establishes sister chromatid cohesion by topologically entrapping two DNA molecules^3,4,17–19^. Cohesion maintains sister loci in proximity to facilitate homology-directed repair of DNA double-strand breaks^20–24^ and provides mechanical stability at centromeres to ensure faithful chromosome segregation in mitosis^1–4,25–28^. Both functions rely on the same cohesin core complex, which associates with distinct regulatory cofactors to promote either loop extrusion or cohesion. While cohesin’s role in shaping elaborate loop architectures is well established (reviewed in ^1,28–31^), it remains unclear how cohesion is positioned relative to loops and boundary organization on replicated chromosomes.

Because loop extrusion and cohesion act on the same replicated chromosomal template, they might influence each other’s positioning. Loop extrusion folds each sister chromatid, but it also promotes sister chromatid resolution and separation^32–34^. Since cohesion requires stable topological entrapment of sister DNAs, extrusion-driven resolution might redistribute cohesive linkages. Conversely, cohesin-mediated linkages can act as barriers to loop extrusion^35–37^, potentially constraining extrusion-driven resolution. Recent studies in mitotic chromosomes indeed suggest functional coupling between loop extrusion and cohesin engaged in cohesion^37,38^. However, mitosis lacks interphase TAD organization and loops are largely formed by condensin-driven extrusion^39,40^, raising the question of how chromosomal context in interphase determines where cohesion can be retained without compromising ongoing loop extrusion.

Here, we combine sister-chromatid– sensitive chromosome conformation mapping with quantitative profiling of cohesin localization, together with acute perturbations of extrusion and cohesion regulators. This integrated strategy allowed us to define the spatial principles that coordinate cohesin’s dual activities across replicated chromosomes, enabling persistent sister linkage while preserving dynamic loop-based folding.

## RESULTS

### Cohesion integrates into a pre-existing 3D genome architecture

To determine how cohesion affects higher-order chromosome structure, we compared genome-wide chromatin contact distributions before and after replication in human cells. Hi-C analysis of G1-synchronized cells (Fig. S1A, C-D) revealed a TAD organization that was very similar to that of published Hi-C data from G2-synchronized cells^33^ (Fig. 1A), consistent with prior work on cell-cycle-phased single-cell Hi-C data^41^. In contrast, published reference maps from mitotic cells^33^ showed a marked loss of TAD organization (Fig. 1A). Genome-wide quantitative analyses of TAD architectures using insulation score^7^ and Adjusted Rand Index (ARI)^42,43^ metrics demonstrated a strong correlation between G1 and G2 but not between G2 and mitosis (Fig. 1B, C). These data indicate that chromatin folding in G1 cells is largely preserved after replication, suggesting that cohesive links form without globally disrupting pre-existing 3D genome organization.

**Figure 1.**
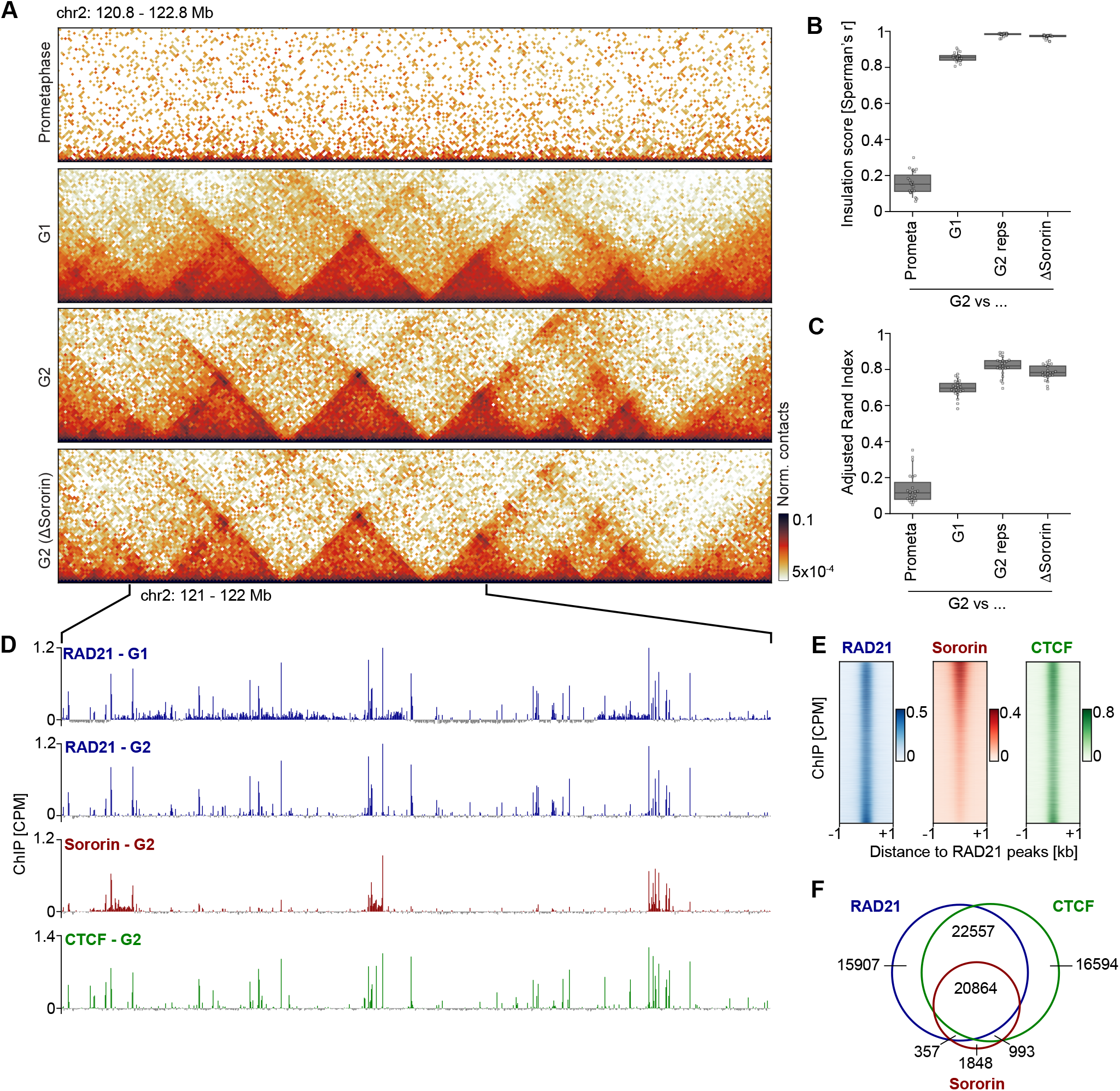
Cohesion integrates into a pre-existing 3D genome landscape. (**A**) Hi-C interaction maps (10 kb resolution, 750 kb separation) from wild-type cells synchronized to prometaphase, G1, or G2 phase, and from G2-synchronized Sororin-depleted (ΔSororin) cells. G1 and G2 maps were downsampled (499 million contacts) to match ΔSororin map; prometaphase map contains 100 million contacts. G1 Hi-C data are merged from n=2 independent experiments. (**B**) Spearman correlation of insulation scores from conditions in (A). Scores were calculated from Hi-C contact maps (20 kb resolution) using a ±500 kb window per chromosome and compared to G2 wild type samples. Biological replicates of wild type G2 (G2 reps) samples served as positive controls: two replicates were merged (333 million contacts), and the third biological replicate was downsampled to match sequencing depth. (**C**) Adjusted Rand Index of TAD annotations (20 kb resolution, 2 Mb maximum separation) calculated per chromosome. Samples were compared as in (B). (**D**) ChIP-Seq signal tracks (1 kb resolution) of RAD21 from G1 cells, and RAD21, Sororin and CTCF from G2 cells. Shown is a zoomed region corresponding to the area in (A). RAD21 G1 and G2 tracks were downsampled to matching sequencing depths (160 million). (**E**) Stacked line profiles of RAD21, Sororin and CTCF ChIP-Seq signal (50 bp resolution) at ±1 kb regions centered on RAD21 peaks identified in G2 cells. Peaks are sorted by the Sororin/RAD21 CPM ratio. (**F**) Venn diagram showing peak overlaps between RAD21, CTCF and Sororin in G2 cells. Reproducible peaks were identified from independent experiments. ChIP-Seq data are CPM-normalized, centered by subtracting the genome-wide mean and represent merged data from n=3 (CTCF) or n=2 (Sororin, RAD21) independent experiments. Wild type G2, Prometaphase and ΔSororin G2 scsHi-C data reanalyzed from^33^.

If TAD structures are preserved after replication, then cohesive cohesin must integrate into an existing chromatin architecture without reshaping it. To test this directly, we analyzed published Hi-C data from G2 cells in which the cohesion factor Sororin^19,44–46^ was depleted using an auxin-inducible degron system^33^. Selective loss of Sororin, and thereby of the cohesive cohesin pool, caused no detectable change in TAD organization relative to wild-type cells (Fig. 1A–C). These findings indicate that the establishment and maintenance of cohesion have minimal structural impact on higher-order chromatin folding.

Since chromatin architecture remains largely unchanged after replication, we next examined how cohesin is positioned along the genome after establishment of cohesion. We performed ChIP–seq for RAD21, a core subunit shared by both loop-extruding and cohesive cohesin complexes^1,2,28^ and compared binding profiles between cells synchronized to G1 or G2 (Fig. S1A-E). RAD21 peaks were highly similar between the two cell cycle phases (Fig. 1D; Fig. S1F), indicating that the establishment of cohesion during S phase does not cause major redistribution of cohesin and instead integrates into a pre-established positional framework.

Following replication, about one third of cohesin complexes mediate cohesion^19,47,48^. To examine how this cohesive pool is distributed along chromosomes, we profiled the distribution of Sororin, a subunit contained in cohesive but not loop-extruding cohesin^19,37,44–46,49^, using ChIP–seq in G2-synchronized cells (Fig. S1B,E). Sororin peaks generally co-localized with RAD21 peaks, yet many RAD21 peaks displayed little or no detectable Sororin enrichment, even at loci with high RAD21 occupancy (Fig. 1D, E, Fig. S1F). Overall, only 36% of RAD21 peaks in G2 overlapped with Sororin peaks (Fig. 1F), consistent with previous studies showing that Sororin-bound cohesin localizes to a distinct subset of cohesin sites^19,46^. These data demonstrate that cohesive cohesin is primarily recruited to existing loop-forming sites rather than to new genomic positions, and that its abundance relative to loop-forming cohesin varies substantially along the genome.

To understand what determines which cohesin sites acquire a cohesive function, we investigated the spatial relationship of RAD21 and Sororin with the major cohesin-targeting protein CCCTC-binding factor (CTCF)^1,12,28–31,50–53^. ChIP-seq of G2-synchronized cells (Fig. S1B, E) showed that although 91% of Sororin peaks colocalize with CTCF, 37% of CTCF peaks contain RAD21 but lack detectable Sororin (Fig. 1D–F; Fig. S1F). These findings suggest that CTCF is necessary but not sufficient for the selective targeting of cohesive cohesin.

Together, these results show that cohesion integrates into the pre-existing cohesin landscape without substantially reshaping overall TAD organization. Notably, cohesive cohesin accumulates at only a subset of loop-forming cohesin sites, revealing a divergent genomic distribution of the two cohesin pools. Because CTCF occupancy alone does not account for this selectivity, these findings raise questions about what other organizational principles might determine where cohesive cohesin is positioned on replicated chromosomes.

### Nested TAD hierarchy defines split and linked loops

To investigate how features of higher-order chromosome folding contribute to the selective enrichment of cohesive cohesin along replicated chromosomes, we examined the relationship between TAD architecture and the distribution of cohesin complexes across larger genomic regions. Comparisons of chromatin contact maps with RAD21 and Sororin ChIP–seq profiles showed that Sororin was primarily enriched at TAD boundaries, whereas many RAD21 peaks inside TAD bodies showed little or no Sororin signal (Fig. 2A, B). Notably, Sororin enrichment was not uniform across boundaries: while RAD21 broadly marked all boundaries (Fig. 2A, B, arrowheads), Sororin concentrated much more prominently at boundaries separating large, high-level domains (Fig. 2A, B, filled arrowheads). In contrast, boundaries delimiting less prominent, nested TADs showed little or no Sororin enrichment (Fig. 2A, B, open arrowheads). These observations suggest that the hierarchy of a boundary within the nested TAD architecture may influence its ability to recruit cohesive cohesin.

**Figure 2.**
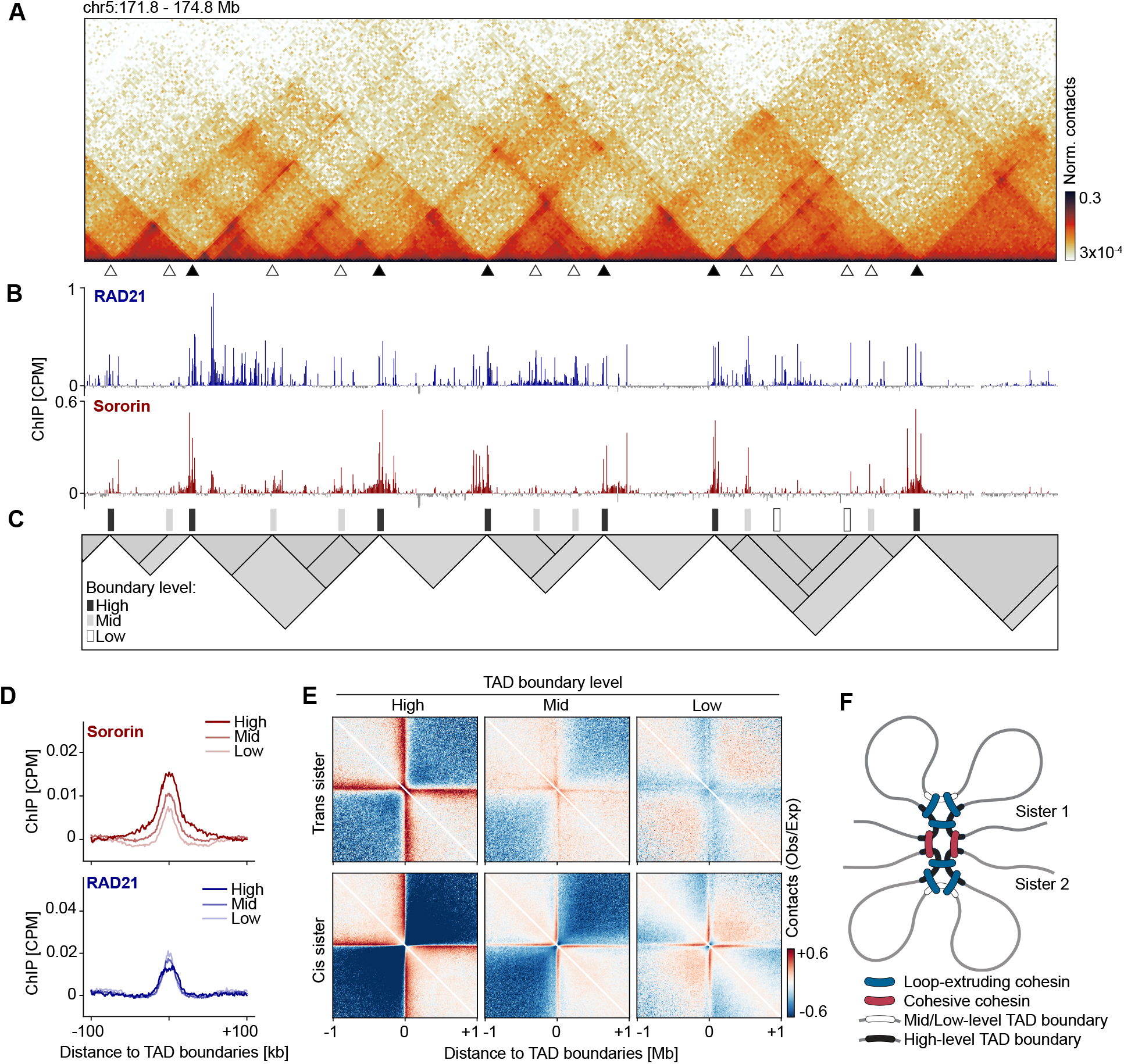
Nested TAD hierarchy defines split and linked loops. (**A**) Representative Hi-C contact map (10 kb resolution, 1.5 Mb separation) from wild type G2 cells. Open arrowheads: TAD boundaries with low/moderate Sororin enrichment; filled arrowheads: TAD boundaries with high Sororin enrichment. (**B**) ChIP-Seq signal tracks (2 kb resolution) of RAD21 (blue) and Sororin (red) at the region shown in (A). (**C**) TAD hierarchy annotation generated by OnTAD. Boundaries are classified into high-, mid- and low-level categories. TAD boundaries were defined at 10 kb resolution and expanded to 20kb for visualization. (**D**) Average profiles of Sororin (upper panel) and RAD21 (lower panel) ChIP-Seq signal (1 kb resolution) at ±100 kb regions centered at high-, mid- and low-level boundaries. (**E**) Average maps of trans sister (upper row) and cis sister (lower row) scsHi-C contacts (10 kb resolution) at ±1 Mb regions centered at high-, mid-, and low-level boundaries, normalized to genome-wide expected contact frequencies. (**F**) Model depicting sister chromatid organization as defined by TAD hierarchy. ChIP-Seq data are CPM-normalized, centered by subtracting the genome-wide mean and represent merged data from n=2 independent experiments. Wild type scsHi-C data reanalyzed from^33^.

To systematically evaluate this relationship, we assigned hierarchy levels to TAD boundaries using OnTAD^43^, which identifies nested domains and stratifies TADs according to their position within the TAD hierarchy (Fig. 2C). Based on the hierarchy of corresponding TADs we classified boundaries into high-, mid-, and low-level categories (Fig. 2C) and analyzed ChIP–seq signals across these groups. Aggregate profiles revealed that Sororin was significantly enriched at high-level boundaries compared to mid- and low-level ones (Fig. 2D, upper panel), whereas RAD21 was more evenly distributed, displaying even modest enrichment at lower-level boundaries (Fig. 2D, lower panel). These data demonstrate, that while loop-forming cohesin broadly associates with boundaries, cohesive cohesin preferentially localizes to the top-level boundaries within the nested TAD hierarchy.

To directly examine how loops and sister-chromatid linkages are organized within the TAD hierarchy, we used sister-chromatid-sensitive Hi-C (scsHi-C). This approach distinguishes contacts within a chromatid (cis sister contacts) from contacts between sister chromatids (trans sister contacts) through 4-thiothymidine labeling of nascent DNA^33,54^. Although previous work reported that trans sister-chromatid contacts are enriched at TAD boundaries, our cohesin ChIP–seq data suggest a more complex distribution within the nested TAD hierarchy. To investigate this, we re-analyzed published scsHi-C data from G2-synchronized cells^33^ to generate aggregate maps centered on TAD boundaries stratified by their position within the hierarchy. Trans sister contacts were enriched in stripes emanating from high-level boundaries, but not at mid-level boundaries, and were even depleted at low-level boundaries relative to the genome-wide average (Fig. 2E, upper row; Fig. S2, upper row). These results indicate that high-level TAD boundaries act as discrete cohesion hubs.

To understand how the variability in trans sister contact density across the boundary hierarchy relates to the loop architectures within sister chromatids, we next analyzed cis sister contact patterns. Average contact maps aligned to TAD boundaries revealed prominent contact enrichment stripes across all hierarchy levels, reflecting local loop formation (Fig. 2E, lower row; Fig. S2, lower row). While these stripes spanned a greater genomic range at high-level boundaries, they remained prominent at mid- and low-level boundaries as well. These results reveal a marked contrast between inter- and intra-sister organization: whereas trans sister contact enrichment is confined to high-level boundaries, cis sister looping occurs at all hierarchical levels. This difference matches the distribution of RAD21 binding observed by ChIP-seq, where cohesin broadly marks boundaries across the hierarchy while cohesive cohesin is restricted to the strongest ones.

Together, these observations show a functional distinction within the TAD hierarchy: mid- and low-level boundaries primarily form loop structures independently on each sister chromatid, whereas high-level boundaries establish loops and at the same time function as specialized cohesion zones (Fig. 2F). This hierarchical TAD organization folds replicated chromosomes into a nested architecture in which small loops form separately on each sister chromatid, while sister chromatids are linked at the anchors of large loops.

### CTCF restricts misalignment of sister chromatids

Given that cohesion is concentrated at high-level TAD boundaries, we asked which molecular factors specify this organization. Because CTCF positions loop-forming cohesin at TAD boundaries^12,50–53,55–57^, we tested whether CTCF also positions cohesion zones. We therefore depleted CTCF in G2-synchronized cells using an auxin-inducible degron system (Fig. S3A-D). To determine how CTCF contributes to the positioning of cohesive cohesin, we profiled Sororin binding by ChIP–seq relative to TAD boundary positions defined in wild-type cells. CTCF depletion strongly reduced Sororin enrichment at TAD boundaries, including high-level boundaries (Fig. 3A–B), whereas Western blotting showed that chromatin-bound RAD21 and Sororin levels remained high (Fig. S3E). Thus, CTCF is dispensable for global chromatin association of cohesive cohesin but required for its focal enrichment at the high-level TAD boundaries that define cohesion zones.

**Figure 3.**
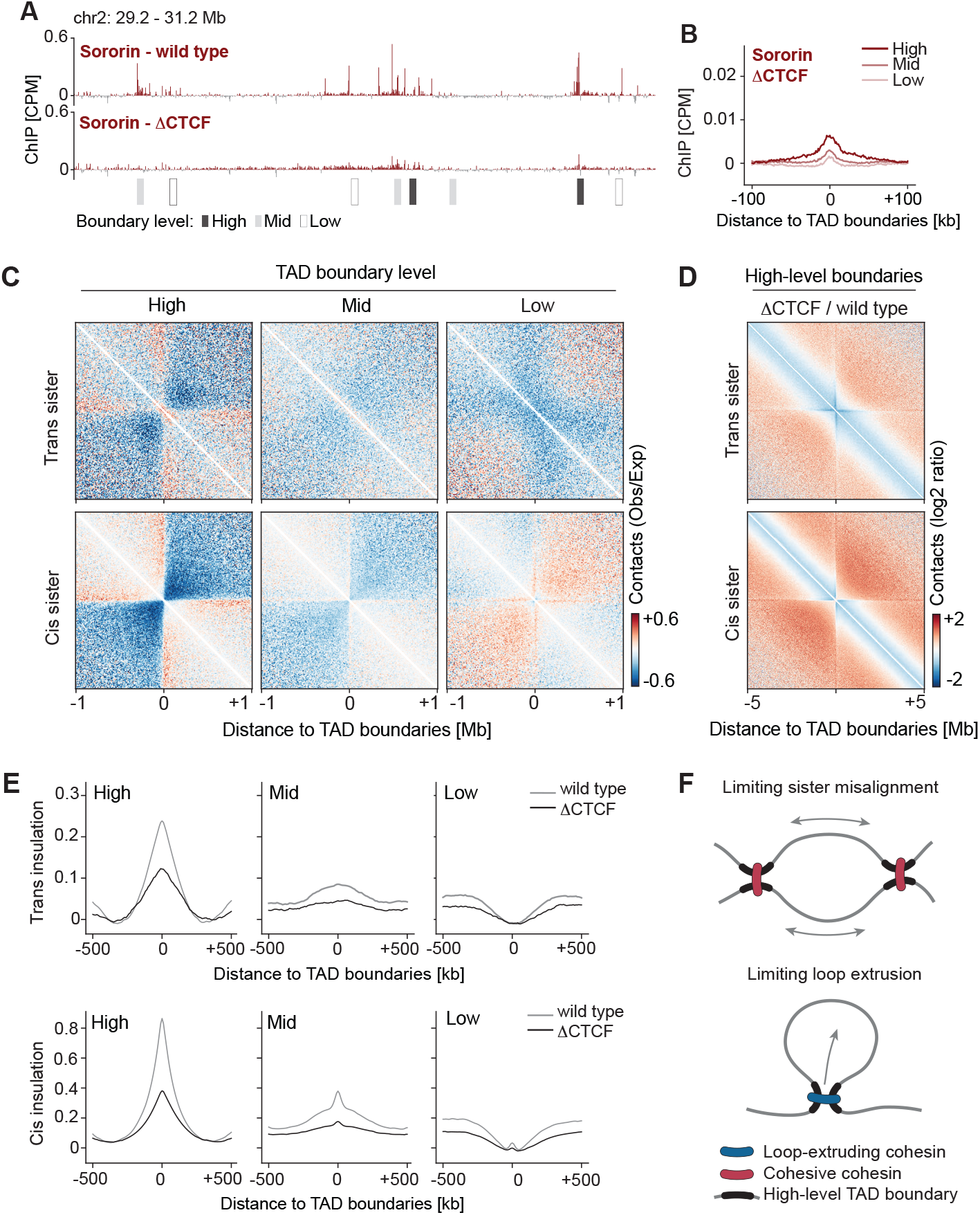
CTCF positions cohesive cohesin and restricts sister chromatid misalignment. (**A**) Representative Sororin ChIP-Seq signal track (2 kb resolution) from G2 wild type and CTCF-depleted (ΔCTCF) cells, with boundaries annotation below. TAD boundaries (defined at 10 kb; expanded to 20 kb for visualization) are indicated. (**B**) Average profiles of Sororin ChIP-Seq signal (1 kb resolution) from G2 ΔCTCF cells at ±100 kb regions centered at high-, mid- and low-level boundaries. (**C**) Average maps of trans sister (upper row) and cis sister (lower row) scsHi-C contacts (10 kb resolution) at ±1Mb regions centered at high-, mid-, and low-level boundaries from G2 ΔCTCF cells, normalized to genome-wide expected contact frequencies. (**D**) Ratio pileups (ΔCTCF/WT) of trans sister (upper panel) and cis sister (lower panel) scsHi-C contacts (20 kb resolution) at ±5Mb regions centered at high-level boundaries. (**E)** Average insulation score profiles calculated from G2 trans sister (upper row) and cis sister (lower row) contacts (10 kb resolution) in wild type (grey line) and ΔCTCF cells (black line) at ±500kb regions centered at high-, mid-, and low-level boundaries. (**F**) Model illustrating CTCF’s role in limiting sister chromatid misalignment and loop extrusion in cis. ChIP-Seq data are CPM-normalized, centered by subtracting the genome-wide mean and represent merged data from n=2 independent experiments. scsHi-C data for ΔCTCF are merged from n=5 independent experiments. Wild type scsHi-C data reanalyzed from^33^.

Given that CTCF is required to concentrate Sororin at TAD boundaries, we next asked how this positioning impacts sister-chromatid organization. Depleting CTCF in G2 phase (Fig. S3F-I) strongly reduced TAD structures in conventional Hi-C maps (Fig. S3J), consistent with CTCF’s established role in boundary positioning of loop-based architectures. We then compared scsHi-C maps from wild-type and CTCF-depleted cells to resolve sister-specific interactions. Upon CTCF depletion, boundary-associated contact stripes were nearly abolished in trans sister maps (Fig. 3C, upper row), indicating that boundary-centered sister–sister organization depends on CTCF. Cis sister maps likewise showed a marked loss of boundary-associated stripes, in line with previous reports^12,53^ (Fig. 3C, lower row). Together, these data show that CTCF not only positions loops within each chromatid but also organizes boundary-associated interactions between sister chromatids.

The displacement of cohesive cohesin away from TAD boundaries in CTCF-depleted cells raised the question of whether this redistribution affects trans sister interactions beyond boundary coordinates. To address this, we calculated ratios of contact densities in CTCF-depleted cells relative to wild-type cells. Aggregate maps of trans sister contacts centered on high-level TAD boundaries revealed a pronounced redistribution from short-range to longer-range interactions at the boundary and at adjacent genomic regions (Fig. 3D, upper panel). Cis sister contacts showed a similar shift toward longer genomic distances (Fig. 3D, lower panel). Genome-wide P(s) curves consistently shifted from short- to long-range contacts in both cis and trans (Fig. S3K). Together, these analyses indicate that CTCF constrains the genomic range of interactions within and between sister chromatids.

The constraint imposed by CTCF on long-range genomic interactions is consistent with its well-established function as a barrier to loop extrusion^12,53,56,58–61^. In conventional Hi-C experiments, such constraints are detected using insulation metrics that quantify contact depletion across domain boundaries. To examine how insulation is organized in replicated chromosomes, we quantified insulation strength across TAD boundaries in cis and trans sister contact maps using insulation score^12^. In wild-type cells, insulation strength scaled with boundary hierarchy: high-level boundaries strongly insulated both cis sister and trans sister contacts, whereas mid- and low-level boundaries showed progressively weaker insulation, with little or no insulation in trans (Fig. 3E, grey lines). CTCF depletion strongly reduced insulation in both cis and trans (Fig. 3E, black lines). Thus, CTCF establishes topological barriers not only for loop interactions within chromatids but also for interactions across sister chromatids, effectively controlling the positioning and the spatial extent of trans sister interactions.

Together, these results establish CTCF as a central regulator of replicated chromosome architecture: by positioning loop-forming cohesin at TAD boundaries and concentrating cohesive cohesin at high-level boundaries CTCF imposes a topological barrier that restricts the genomic range of cis and trans sister interactions (Fig. 3F). By limiting the genomic span of trans sister interactions, CTCF constrains large-scale sister chromatid misalignment while preserving dynamic loop architectures within intervening domains.

### Loop extrusion positions cohesive cohesin within the TAD hierarchy

While our results show that CTCF positions both loop-forming and cohesive cohesin at TAD boundaries, it remains unclear how the two cohesin pools become differentially enriched across the nested TAD hierarchy. One possibility is that this hierarchy-dependent distribution reflects dynamic interactions between the two pools. Loop-forming cohesin is positioned at boundaries through its intrinsic ATP-dependent motor activity^5,6^, enabling translocation along chromatin until extrusion is stalled at CTCF boundary elements^12,53,56,57,59,62,63^. In contrast, cohesive cohesin is not known to have a motor activity and it does not contain the NIPBL subunit required for ATP hydrolysis and DNA translocation^5,6,64–74^. Its genomic distribution may therefore depend on forces generated by loop-extruding cohesin. Indeed, mitotic studies suggest that loop-extruding cohesin can push cohesive cohesin along DNA toward boundary elements^37^.

However, how such extrusion-driven pushing would generate the pronounced TAD hierarchy–dependent enrichment of cohesive cohesin observed in interphase remains unclear.

To test whether loop extrusion is required to concentrate cohesive cohesin at high-level boundaries in interphase, we depleted NIPBL in G2-synchronized cells using an auxin-inducible degron (Fig. S4 A-D), suppressing loop extrusion while preserving established cohesion. Hi-C analysis of published data^34^ showed a marked loss of TAD structures under these conditions (Fig. S4E), whereas chromatin-bound Sororin levels remained high (Fig. S4F). ChIP–seq profiling revealed that Sororin largely lost its enrichment at high-level TAD boundaries and instead became broadly distributed across the genome (Fig. 4A, red tracks, 4B, upper panel). RAD21 ChIP–seq likewise showed reduced boundary-centered enrichment and a more uniform genomic distribution (Fig. 4A, blue tracks, 4B, lower panel). Thus, loop extrusion is required to concentrate cohesive cohesin at high-level boundaries.

**Figure 4.**
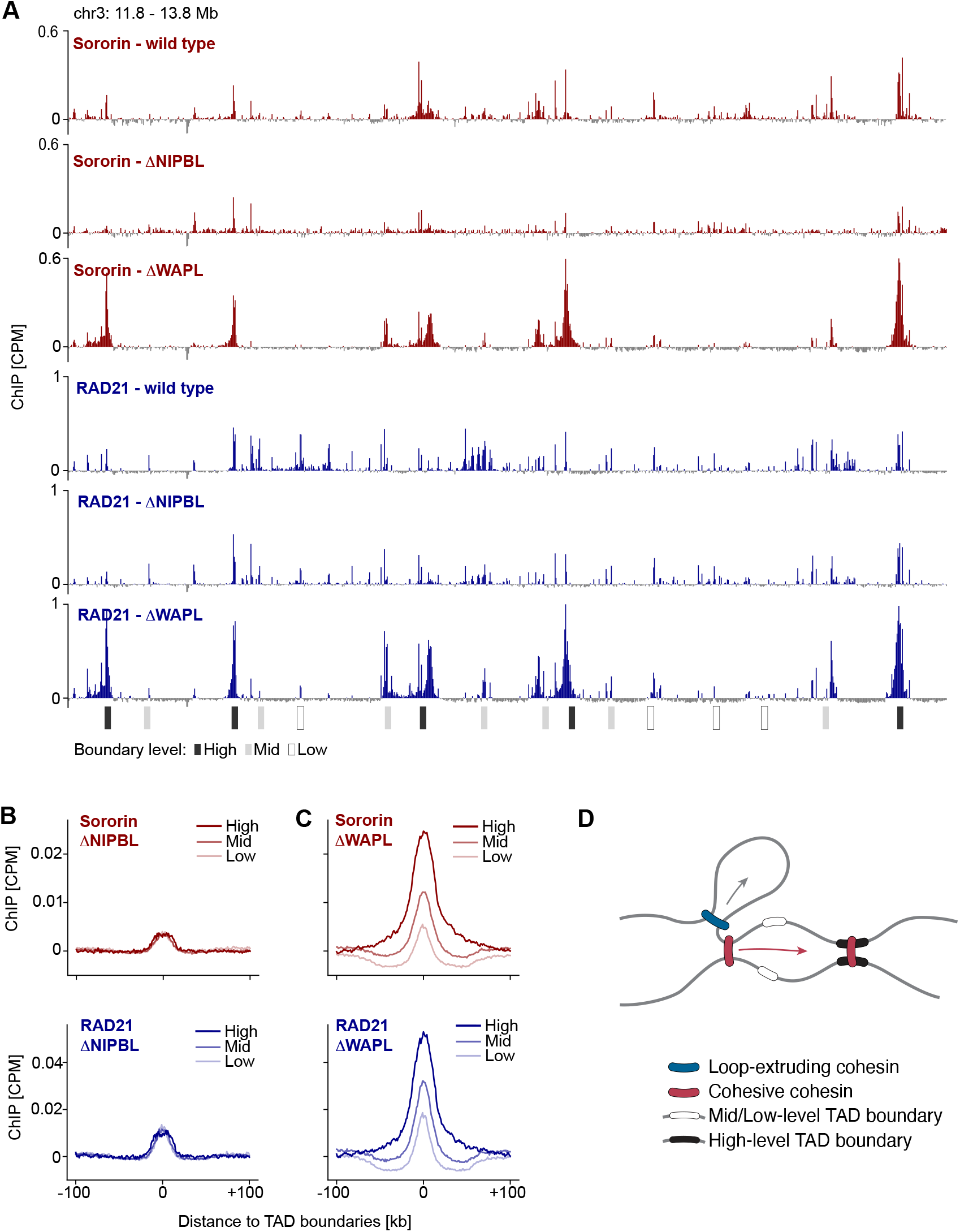
Loop extrusion concentrates cohesive cohesin at high-level TAD boundaries. (**A**)Representative ChIP-Seq tracks (2 kb resolution) of Sororin (red) and RAD21 (blue) from G2 wild type, NIPBL-depleted (ΔNIPBL) and WAPL-depleted (ΔWAPL) cells, with boundaries annotation below. TAD boundaries (defined at 10 kb; expanded to 20 kb for visualization) are indicated. (**B**) Average profiles of Sororin (upper panel, red) and RAD21 (lower panel, blue) ChIP-Seq signal (1 kb resolution) from G2 ΔNIPBL cells at ±100 kb regions centered at high-, mid- and low-level boundaries. (**C**) Average profiles of Sororin (upper panel, red) and RAD21 (lower panel, blue) ChIP-Seq signal (1 kb resolution) from G2 ΔWAPL cells at ±100 kb regions centered at high-, mid- and low-level boundaries. (**D**) Model depicting loop extrusion-mediated positioning of cohesive cohesin at strong TAD boundaries. ChIP-Seq data are CPM-normalized, centered by subtracting the genome-wide mean and represent merged data from n=2 independent experiments.

If loop extrusion promotes the accumulation of cohesive cohesin at high-level boundaries, then increasing the extrusion processivity should amplify this effect. To test this, we depleted WAPL in G2-synchronized cells using a degradation tag system (Fig. S5A-E), which increases cohesin residence time and promotes hyper-processive extrusion^12,13,75–80^. Hi-C maps showed stronger corner signals and increased long-range interactions (Fig. S5F), consistent with enhanced extrusion, whereas chromatin-bound RAD21 and Sororin levels remained high (Fig. S5G). ChIP–seq profiling showed that Sororin strongly concentrated at high-level TAD boundaries, reaching levels substantially higher than in wild-type cells, while it decreased at low-level boundaries (Fig. 4A red tracks, 4C, upper panel). RAD21 displayed a similar redistribution (Fig. 4A, blue tracks, 4C, lower panel). Thus, hyper-processive extrusion amplifies focal accumulation of both cohesive and loop-forming cohesin at high-level boundaries.

Together, these findings demonstrate that loop extrusion dynamics play a central role in the positioning of cohesive cohesin (Fig. 4D). Our data are consistent with a model in which loop-extruding cohesin can push cohesive cohesin toward boundary elements^37^, but further show that extrusion does not uniformly concentrate cohesive cohesin at all boundaries in interphase cells. Instead, loop extrusion can either increase or decrease cohesive cohesin enrichment at boundaries, depending on the interplay between boundary hierarchy and extrusion processivity. Extrusion dynamics thus provide a mechanism for distributing cohesive cohesin within the nested TAD architecture.

### Loop extrusion shapes cohesion zones

Because loop-extrusion processivity affects the positioning of both cohesin pools, we next asked how altering extrusion dynamics affects the nested architecture of linked and split loops in replicated chromosomes. To directly relate cohesin dynamics to chromosome conformation, we used scsHi-C to quantify both cis sister loops and trans sister contacts in G2 cells under conditions in which loop extrusion is either suppressed by NIPBL depletion or rendered hyper-processive by WAPL depletion.

To determine how loop extrusion contributes to the formation of cohesion zones, we compared published scsHi-C maps from G2-synchronized wild-type cells^33^ with those from cells in which NIPBL was depleted after DNA replication^34^. The prominent contact stripes emanating from high-level boundaries in both cis and trans sister contacts maps were almost completely lost (Fig. 5A). Contact-density ratios relative to wild type at high-level boundaries confirmed a marked depletion of boundary-centered enrichment and a redistribution of trans sister contacts toward shorter genomic distances (Fig. 5B). Thus, consistent with the dispersal of cohesive cohesin observed by ChIP–seq in NIPBL-depleted cells, inhibition of loop extrusion abolishes the formation of defined cohesion zones at high-level TAD boundaries.

**Figure 5.**
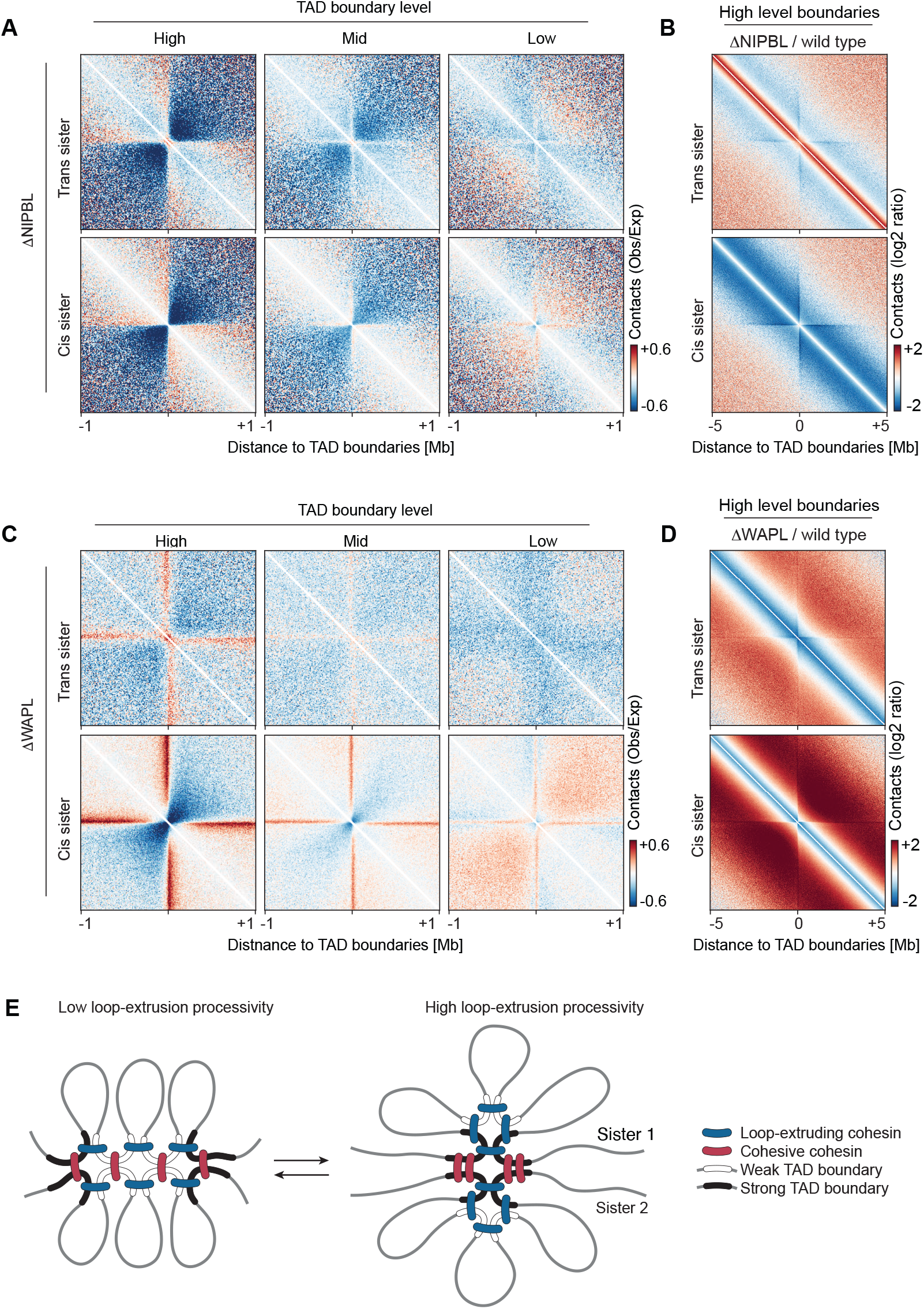
Loop extrusion positions trans sister contacts at high-level boundaries. (A) Average maps of trans sister (upper row) and cis sister (lower row) scsHi-C contacts (10 kb resolution) at ±1Mb regions centered at high-, mid-, and low-level boundaries from G2 cells upon NIPBL depletion (ΔNIPBL), normalized to genome-wide expected contact frequencies. (**B**) Ratio pileups (ΔNIPBL/wild type) of trans sister (upper panel) and cis sister (lower panel) scsHi-C contacts (20 kb resolution) at ±5Mb regions centered at high-level boundaries. (**C**) Average pileups of trans sister (upper row) and cis sister (lower row) scsHi-C contacts (10 kb resolution) at ±1Mb regions centered at high-, mid-, and low-level boundaries from G2 cells upon WAPL depletion (ΔWAPL), normalized to genome-wide expected contact frequencies. (**D**) Ratio pileups (ΔWAPL/wild type) of trans sister (upper panel) and cis sister (lower panel) scsHi-C contacts (20 kb resolution) at ±5Mb regions centered at high-level boundaries. (**E**) Model illustrating the role of loop extrusion processivity in positioning cohesive cohesin within the TAD hierarchy. scsHi-C data for ΔWAPL are merged from n=2 independent experiments. Wild type scsHi-C data reanalyzed from^33^. ΔNIPBL scsHi-C data reanalyzed from^34^.

We next asked how hyper-processive loop extrusion reshapes boundary-associated organization. We performed scsHi-C in WAPL-depleted G2 cells and observed a pronounced reorganization of boundary features. In cis, contact stripes remained prominent, but insulation between adjacent domains was reduced (Fig. 5C, lower row), consistent with previous observations^12,81,82^. In trans, stripes at high-level boundaries remained visible but did not increase in strength compared to wild-type cells (Fig. 5C, upper row), despite elevated levels of Sororin-bound cohesin (Fig. 4A, C). Instead, contact-density ratios relative to wild type at high-level boundaries revealed a genome-wide shift toward longer-range trans sister contacts at boundaries and in adjacent regions (Fig. 5D), consistent with increased separation and reduced local alignment of sister chromatids. Together, these patterns suggest that under hyper-processive extrusion, high-level boundaries interact with a broader, less focused region on the sister chromatid.

Together, these perturbations show that loop extrusion dynamics fine-tune the nested architecture of replicated chromosomes (Fig. 5E). When extrusion is inhibited, trans-sister interactions are no longer focused at high-level boundaries; when extrusion is hyper-processive, cis loops expand and trans-sister contacts extend to longer genomic distances. These results support a model in which balanced extrusion dynamics promote nested cis loop structures that form independently on each sister chromatid, while global sister chromatid alignment is maintained through discrete cohesion zones at high-level TAD boundaries.

### Clusters of CTCF sites define broad cohesion zones

High-level boundaries insulate stronger both in cis and trans (Fig. 3E), and strong TAD boundaries often contain multiple closely spaced CTCF sites^60,83,84^. Consistent with this, we found that 49% of the high-level TAD boundaries contain three or more CTCF peaks (Fig. S6A), in contrast to 36% and 25% of medium-and low-level boundaries, respectively. This raised the question of how cohesive cohesin is positioned relative to internal boundary organization. To address this, we analyzed the distribution of RAD21 and Sororin around CTCF sites located within high-level TAD boundaries compared with CTCF sites at other genomic regions.

Inspection of representative loci showed that Sororin, like RAD21, formed sharp peaks centered on CTCF peaks (Fig. 6A; Fig. S6B). Within high-level boundary regions, however, Sororin signal remained elevated between neighboring CTCF sites compared to the genome-wide average. This suggests that cohesive cohesin is retained not only at individual CTCF sites but also across the intervening boundary region.

**Figure 6.**
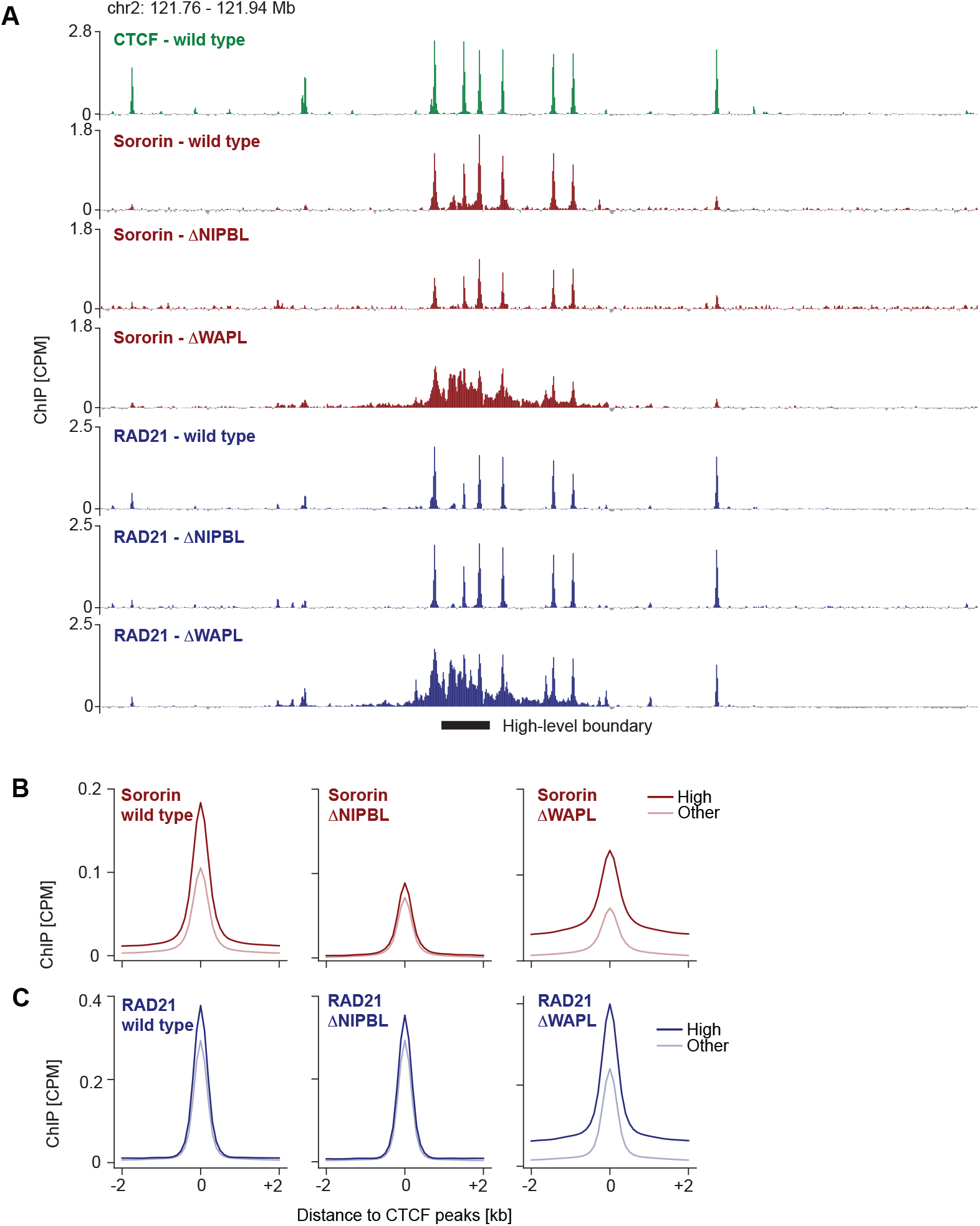
Clusters of CTCF sites define broad cohesion zones. (A) Representative ChIP-Seq tracks (200 bp resolution) of CTCF (green) from G2 wild type, and Sororin (red) and RAD21 (blue) from G2 wild type, NIPBL-depleted (ΔNIPBL) and WAPL-depleted (ΔWAPL) cells. Position of a high-level boundary (10 kb long) is indicated below. (**B**) Average profile of Sororin ChIP-Seq signal (100 bp resolution) from G2 cells under wild type (left panel), ΔNIPBL (middle panel) and ΔWAPL (right panel) conditions at ±2 kb regions centered at CTCF peaks within high-level boundaries (High) (n= 8034 peaks) or elsewhere in the genome (Other) (n=53351 peaks). (**C**) Average profile of RAD21 ChIP-Seq signal (100 bp resolution) from G2 cells under wild type (left panel), ΔNIPBL (middle panel) and ΔWAPL (right panel) conditions at ±2 kb regions window centered at CTCF peaks within high-level boundaries (High) (n=8034 peaks) or elsewhere in the genome (Other) (n=53351 peaks). ChIP-Seq data are CPM-normalized, centered by subtracting the genome-wide mean and represent merged data from n=3 (CTCF) or n=2 (Sororin, RAD21) independent experiments.

To test whether this pattern is a general feature of high-level boundaries, we performed aggregate analyses across all CTCF peaks. At CTCF peaks within high-level boundary regions, Sororin displayed both increased peak amplitude and an elevated inter-peak baseline compared with CTCF peaks elsewhere in the genome (Fig. 6B, left panel), whereas RAD21 showed only a modest elevation of inter-peak signal (Fig. 6C, left panel). To assess whether the elevated baseline could be explained by contributions from neighboring CTCF peaks, we analyzed Sororin occupancy at CTCF peaks without other peaks in their close vicinity (±2.5 kb). Baseline Sororin signal remained elevated at high-level boundaries compared to elsewhere in the genome (Fig. S6C), indicating that the pronounced inter-peak enrichment of Sororin is not only attributable to signal from adjacent CTCF peaks. Thus, high-level TAD boundary regions form extended cohesion zones in which cohesive cohesin accumulates both at CTCF peaks and across the intervening boundary interval.

Since loop extrusion positions cohesive cohesin to boundaries, it may also regulate the fine scale distribution of cohesive cohesin around CTCF sites. When we analyzed Sororin binding in NIPBL-depleted cells, Sororin signal was markedly reduced at CTCF peaks within boundary regions, and the elevated inter-peak baseline observed in wild-type cells almost completely disappeared (Fig. 6A, red tracks, 6B, middle panel). Thus, loop extrusion is required to concentrate cohesive cohesin not only at CTCF sites within TAD boundaries but also across the intervening regions that define extended cohesion zones.

To assess how hyper-processive loop extrusion reshapes the fine scale boundary organization, we examined the effects of WAPL depletion on cohesin distribution. ChIP–seq profiles revealed a striking redistribution of Sororin within high-level TAD boundaries: Sororin signal at CTCF peaks decreased relative to genome-wide average, both within boundary regions and across the genome, while the inter-peak baseline within boundary regions became strongly elevated (Fig. 6A, red tracks, 6B, right panel). RAD21 showed a similar accumulation between neighboring CTCF sites at high-level boundaries (Fig. 6A, blue tracks, 6C, right panel). These results indicate that hyper-processive extrusion reshapes cohesion zones by shifting both cohesin pools from sharp, CTCF-centered peaks into broader inter-peak regions within high-level TAD boundaries.

Together, these results show that clustering of CTCF sites at high-level boundaries contributes to the establishment of broad cohesion zones, where loop extrusion shapes the fine-scale distribution of cohesive cohesin: under wild type conditions, Sororin is enriched both at CTCF peaks and across the intervening boundary interval, whereas hyper-processive extrusion shifts Sororin from individual peaks into the inter-peak regions. Thus, loop extrusion dynamics actively distribute cohesive cohesin across CTCF clusters, transforming discrete binding sites into continuous cohesion zones.

## DISCUSSION

The replicated interphase genome faces a fundamental organizational challenge: it must keep sister chromatids in proximity to enable homology-directed DNA repair and faithful chromosome segregation while maintaining the dynamic loop structures that support gene regulation. Both requirements are mediated by cohesin, yet cohesin’s two principal activities are inherently antagonistic: loop extrusion promotes chromatid separation, whereas cohesion resists separation and can impede loop extrusion^2,32–37^. Our findings reconcile this paradox by revealing a spatial partitioning of cohesin function on replicated chromosomes. Rather than co-localizing uniformly with loop-extruding cohesin, cohesive cohesin concentrates at discrete genomic positions defined by the nested TAD hierarchy. This segmentation creates cohesion zones that maintain large-scale sister chromatid alignment, while loop extrusion predominates within the intervening chromatin. Thus, hierarchical boundary organization enables mechanical pairing and dynamic folding to coexist on the same replicated chromosome template.

### Interplay of cohesin and CTCF defines cohesion zones

Our data are consistent with a model in which loop extrusion moves cohesive cohesin toward boundary elements, as proposed from studies in mitotic cells ^37^. In interphase, however, the two cohesin pools diverge in their distributions, raising the question of how cohesive cohesin becomes selectively enriched at high-level boundaries. This divergence may reflect dynamic interactions between cohesin and CTCF. Individual CTCF peaks act as semi-permeable barriers that transiently stall cohesin yet can be bypassed^56,58,59,85^. In contrast, high-level boundary regions containing multiple CTCF peaks establish more effective barriers, maintaining insulation even upon occasional bypass of individual CTCF peaks^60,83,86^. In this context, extrusion can enrich cohesin at boundary regions but can also redistribute cohesin away from individual CTCF peaks, depending on extrusion dynamics and local boundary context. Loop extrusion processivity then dictates the degree to which cohesive cohesin is biased toward strong boundaries versus weak ones. Notably, loop-extruding cohesin dissociates within minutes and reforms loops elsewhere in the genome, whereas cohesive cohesin persists on chromatin for many hours throughout interphase^46–48^. This distinct chromatin binding time provides a simple mechanism for selective accumulation: cohesive cohesin accumulates at high-level boundaries over time, whereas the loop-extruding pool remains more evenly distributed across boundaries of different hierarchy levels. Barrier permeability, loop extrusion processivity, and differential cohesin turnover on chromatin thereby give rise to distinct distributions of loops and cohesive linkages within the same CTCF-defined positional framework.

### A shared 3D architecture for gene regulation and DNA repair

The formation of cohesion zones on replicated chromosomes provides a structural framework for homology-directed DNA repair. By limiting the genomic range of trans–sister contacts, cohesion zones confine homology search to regions on the order of a high-level TAD. Such spatially focused sampling offers a major kinetic advantage for identifying homologous repair templates compared with a genome-wide search^2,23,87–90^. At the same time, the sparseness of cohesion zones along chromosome arms suggests why DNA double-strand breaks trigger local architectural remodeling to further enhance search efficiency, by increasing local looping and by forming a cohesive clamp near the break^23,88,91,92^. Cohesin thereby effectively converts a high-level TAD into a homology-sampling unit, enabling rapid and accurate repair.

Restricting cohesion to defined zones may also minimize the extent to which cohesive cohesin impedes loop extrusion^35–37^, leaving most of the chromatin fiber permissive for extrusion-driven looping that supports enhancer–promoter communication^14–16^. The nested TAD hierarchy therefore separates functions across scales: high-level boundaries maintain sister alignment, whereas intervening domains maintain dynamic cis loop architectures on each chromatid. By integrating these distinct modes of cohesin activity within the TAD hierarchy, replicated chromosomes can support homology-directed DNA repair and transcriptional regulation within the same 3D architecture.

### Conclusions and future directions

Our findings raise important questions about the organization of replicated chromosomes. First, where and how is cohesion initially established during S phase, and how do these sites relate to the steady-state accumulation of cohesive cohesin in G2? Second, which features, besides CTCF sites, define where cohesion-zones form, including chromatin state, transcriptional context, and additional architectural factors? Third, to what extent is cohesive cohesin positioning influenced by forces other than cohesin-mediated loop-extrusion, for example by other SMC complexes such as SMC5/6^1,29,93^, by replication-associated machineries such as MCM ^94^, or by the transcription machinery^52,95–100^? Our sister-specific conformation capture combined with protein localization and degron-based perturbations provides a powerful framework to address these and other questions emerging from our study.

Our study highlights a general design principle by which SMC-regulated chromosome architectures can be tuned by balancing processivity, residence time, boundary strength, and topological linkage. In mitosis, condensin I and II generate nested loop architectures through differential processivity and residence time^40,101–103^, and the removal of CTCF-defined boundaries coincides with a major reconfiguration of chromosome folding^33,39,41,104^. In developing B cells, reduced expression of WAPL increases cohesin loop-extrusion processivity to enable long-range V(D)J recombination^105–107^. In replicated chromosomes, we find that an interplay between cohesin pools with distinct kinetics establishes discrete cohesion zones that tether sister chromatids while preserving a dynamic landscape of regulatory loops. Together, these findings suggest that a single positional framework, encoded by the genomic positions of CTCF binding sites, can generate fundamentally different chromosome architectures across the cell cycle. By tuning extrusion dynamics, boundary strength, and topological linkage, this framework supports diverse functions such as transcription, DNA repair, and chromosome segregation.

## ACKNOWLEDGMENTS

We thank the IMBA/IMP/GMI BioOptics and Molecular Biology Service and the Vienna BioCenter Next Generation Sequencing facilities for technical support, J.-M. Peters for providing CTCF-AID and CTCF-AID/SCC1-Halo HeLa cell lines, S. Schüchner and E. Ogris for the generation of mouse anti-Sororin antibody, and I. Patten for comments on the manuscript. The computational results presented were obtained using the CLIP cluster.

Research in the laboratory of DWG has been supported by the Austrian Academy of Sciences, the Vienna Science and Technology Fund (WWTF; project LS19-001), and the European Research Council (ERC) under the European Union’s Horizon 2020 research and innovation programme (grant agreement no. 101019039). DWG is also an adjunct professor at the Medical University of Vienna. ZT has received a Hertha Firnberg Programme fellowship of the Austrian Science Fund (FWF T 1246). SK has received a Boehringer Ingelheim Fonds PhD fellowship.

Research in the laboratory of AG has been supported by the Austrian Academy of Sciences, the Austrian Science Fund (FWF; project SFB F 8804-B “Meiosis”), and the European Research Council (ERC) under the European Union’s Horizon 2020 research and innovation programme (grant agreement no. 101163751).

## Author contributions

Conceptualization: ZT, DM, AG, DWG, CCHL

Methodology: ZT, DM

Software: DM

Formal analysis: DM, ZT

Investigation: ZT, DM, SK

Visualization: DM, ZT

Data curation: CCHL, DM, ZT

Funding acquisition: DWG, ZT

Supervision: DWG, AG

Writing – original draft: DWG

Writing – review & editing: All authors

## Declaration of generative AI and AI-assisted technologies in the manuscript preparation process

ChatGPT (version 5.2, OpenAI) was used to assist in improving the grammar of the manuscript text. No content was generated or modified relating to the scientific findings, data interpretation, or conclusions. All edits were reviewed and approved by the authors.

## METHODS

### Cell culture

HeLa Kyoto cells were provided by S. Narumiya (Kyoto University) and validated via Multiplex Human Cell line Authentication (MCA). Cells were maintained in high-glucose DMEM (Capricorn Scientific, DMEM-HPA-P10) supplemented with 10% (v/v) FBS (Gibco, 10270-106), 1% (v/v) GlutaMAX (Invitrogen, 35050038), and 1% (v/v) penicillin-streptomycin (Sigma-Aldrich, P0781), hereafter referred to as ‘growth medium’. NIPBL-AID and WAPL-dTAG lines were maintained under selection with 6 mg/ml blasticidin S (Thermo Fisher Scientific, A1113903). All cultures were grown at 37°C in a humidified incubator with 5% CO_2_ and routinely tested for mycoplasma. All cell lines used in this study are listed in Table S1.

### Cell synchronization and targeted protein degradation

For synchronization at the G1/S boundary, asynchronous cells were treated with 2 mM thymidine (Sigma-Aldrich, T1895) for 18 h, washed twice with pre-warmed growth medium, and released for 8 h. Cells were then re-arrested with 3 mg/ml aphidicolin (Sigma-Aldrich, A0781) for 16 h. G1/S populations were harvested immediately following this block. For synchronization in G2 phase, cells were released from aphidicolin into fresh growth medium for 4 h and subsequently treated with 9 mM RO-3306 (Sigma-Aldrich, SML0569) for 20 h prior to harvest.

To induce targeted protein depletion, degron ligands were added to the medium as follows:

- NIPBL-AID: 1 mM 5-Ph-indole-3-acetic-acid (5-Ph-IAA; Bio Academia, 30-003) for 16 h, starting 8 h post-aphidicolin release.
- WAPL-dTAG: 1 mM dTAG-7 (Tocris, 6912) for 16 h, starting 8 h post-aphidicolin release.
- CTCF-AID/SCC1-HALO (scsHiC): 500 mM indole-3-acetic acid (IAA; Sigma-Aldrich, I5148) for 12 h, starting 12 h post-aphidicolin release.

-CTCF-AID (ChIP-Seq/Chromatin fractionation): 500 mM IAA (Sigma-Aldrich, I5148) for 16 h, starting 8 h post-aphidicolin release.

For sister-chromatid-sensitive Hi-C (scsHi-C) labeling, 2 mM 4-thio-thymidine (4sT; Carbosynth, NT06341) was added at the onset of the aphidicolin block and maintained until harvest.

All cell lines used in this study are listed in Table S1.

### Flow cytometry and cell cycle analysis

Cell cycle distribution was determined by quantifying DNA content via propidium iodide (PI) staining and Histone H3 Serine 10 phosphorylation (pH3S10). Unless otherwise stated, all centrifugation steps were performed at 1,100 x g for 1 min. Harvested cells (2 × 10_5_) were washed with PBS and fixed in 70% ethanol (Sigma-Aldrich, 32221) for at least 1 h at 4°C. Fixed cells were permeabilized with 0.25% Triton X-100 (Sigma-Aldrich, 327371000) for 15 min on ice and subsequently incubated with 0.25 mg anti-phospho-H3 (Ser10) antibody (Millipore, 05-806) for 1 h at room temperature (RT). After washing with 1% BSA in PBS, cells were incubated with Alexa Fluor 488-conjugated anti-mouse IgG (1:300; Molecular Probes, A11001) for 30 min at RT. Cells were washed and resuspended in PBS containing PI (50 mg/ml; Sigma-Aldrich, 81845) and RNAse A (200 mg/ml; Qiagen, 19101) for 30 min at RT. Data were acquired on Penteon flow cytometer using NovoCyte software. The PI signal was detected using a B615 (615/20) filter, and Alexa Fluor 488 was detected using a B525 (525/45) filter. Data analysis was performed using FlowJo v.10. The gating strategy consisted of: (1) identification of main cell population using FSC-A/SSC-A, (2) doublet exclusion using FSC-A/FSC-H and PI-A/PI-H, (3) quantification of cell cycle phases by plotting pH3S10) intensity (y-axis) against DNA content (x-axis).

### Chromatin fractionation

Synchronized cells (up 2 × 10_6_) were harvested by trypsinization in the presence of 9 mM RO-3306 and appropriate protein degron ligands to maintain cell cycle state and protein depletion. Cells were washed in PBS, collected by centrifugation (2,000 x g, 3 min, 4°C), and stored at -70°C. Pellets were resuspended in ice-cold lysis buffer (20 mM Tris-HCl pH 7.5, 100 mM NaCl, 5 mM MgCl2, 10% glycerol, 0.2% NP-40 1 mM DTT and 1X protease inhibitor cocktail (Roche, 11836170001)) and incubated for 20 min on ice. The soluble fraction was collected following centrifugation (2,000 x g, 3 min, 4°C). The remaining chromatin pellet was washed three times with ice-cold lysis buffer and then solubilized by incubation with Benzonase (1:500; Sigma-Aldrich, 70746) in lysis buffer for 30 min at room temperature. The remaining insoluble fraction was removed by centrifugation (16,000 x g, 20 min, 4°C), and the supernatant was collected as the chromatin-bound fraction.

### Western blotting

Protein samples were resolved on NuPAGE 4-12% Bis-Tris (Thermo Fisher Scientific, NP0335BOX) or, specifically for NIPBL-AID detection, 3-8% Tris-Acetate (Thermo Fisher Scientific, EA0378BOX) polyacrylamide gels. Proteins were transferred to 0.45 µm PVDF membranes (Amersham Hybond P; Sigma-Aldrich, GE10600023) in transfer buffer containing 20% ethanol, or without ethanol for NIPBL-AID detection. Membranes were blocked in 5% non-fat milk in PBS containing 0.05% Tween-20 (PBST) and incubated overnight at 4°C with the following primary antibodies: anti-CTCF (1:1000, Cell Signaling Technology, 2899), anti-MAU2 (1:1000, Abcam, ab183033), anti-NIPBL (1:800, Absea, 010702F01), anti-WAPL (1:1000, Santa-Cruz Biotechnology, sc-365189), anti-RAD21 (1:1000, Abcam, ab992 or ab217678) and anti-Sororin (1:50, in house; as described in _34_). Loading controls were detected using antibodies against GAPDH (1:2500, Abcam, ab9485), a-Tubulin (1:1000, Abcam, ab52866), Vinculin (1:10 000, Abcam, ab129002) and Histone H3 (1:1000, Cell Signaling Technology, 4499). Membranes were washed three times in PBS for 10 min each and subsequently incubated with HRP-conjugated secondary antibodies: goat anti-rabbit (1:5000, Bio-Rad, 1706515), goat anti-mouse (1:5000, Bio-Rad, 1706516) and goat anti-rat (1:5000, Amersham, NA935). Membranes were washed three times in PBS for 10 min each. Chemiluminescent signals were developed using Clarity Max ECL substrate (Bio-Rad, 1705062) and visualized on a ChemiDoc imaging system (BioRad). All antibodies used in this study are listed in Table S2.

### Hi-C and scsHiC sample preparation

Synchronized cells (2 × 10_6_) were harvested by trypsinization in the presence of appropriate compounds to maintain cell cycle phase and protein depletion. Cells were washed with PBS, fixed in 1% formaldehyde (Thermo Fisher Scientific, 28906) for 4 min at room temperature (RT) and quenched with 20 mM Tris (pH 7.5). Pellets were collected and stored at -70°C. Unless otherwise stated, all centrifugation steps were performed at 2,500 x g for 5 min. Cells were incubated in ice-cold lysis buffer (10 mM Tris-HCl pH 8.0, 10 mM NaCl, 0.2% NP-40, 1x protease inhibitor cocktail (Roche, 11836170001)) for 30 min at 4°C. Following lysis, nuclei were collected by centrifugation and washed once with 1x ice-cold digestion buffer (NEB, B0543). Nuclei were permeabilized in digestion buffer containing 0.1% SDS for 10 min at 65°C with shaking (300 rpm). SDS was quenched by the addition of 1% Triton-X for 10 min on ice. Chromatin was digested overnight at 37°C with 375 U DpnII (NEB, R0543) at 800 rpm, followed by heat inactivation at 65°C for 20 min. DNA ends were biotinylated using 50 U Klenow polymerase (NEB, M0210) in 1x NEB2 Buffer (NEB, B7002) supplemented with 38 µM each of biotin-dATP, dCTP, dGTP and dTTP for 1 h at 37°C (800 rpm). DNA fragments were ligated for 4 h at RT in a reaction containing 1x T4 DNA ligase buffer (Thermo Fisher Scientific, B69), 0.1% Triton-X, 100 µg/ml BSA, 50 U T4 DNA ligase (Thermo Fisher Scientific, EL0011). Crosslinks were reversed overnight at 65°C (Hi-C) or for 6 h at 65°C (scsHi-C). DNA was purified using the DNeasy Blood & Tissue Kit (Qiagen, 69506) and sheared to a target size of ∼500 bp using a Covaris S2 sonicator (duration: 25 sec, duty cycle: 10%, intensity: 5.0, cycles/burst: 200). Following size selection with AMPure XP beads (Beckham Coulter, A63881), biotinylated fragments were captured using Streptavidin C1 beads (Invitrogen, 65001) for 60 min at RT with rotation. For scsHi-C, an additional thiol-conversion step was performed using 0.45 mM OsO_4_ (Sigma-Aldrich, 75632) and 180 mM NH_4_Cl (pH 8.88) for 3 h at 60°C, followed by ethanol precipitation. Libraries were prepared using NEBNext Ultra II DNA Library Prep Kit (NEB, E7645). DNA was eluted from streptavidin beads using 95% formamide (Sigma-Aldrich, F9037) in 10 mM EDTA (pH 8.0) for 2 min at 65°C. Following ethanol precipitation, libraries were amplified according to the NEBNext Ultra II protocol and purified with AMPure XP beads (0.55x ratio). Libraries were sequenced on Illumina NextSeq2000, NovaSeq SP or NovaSeqX platforms.

### ChIP-Seq

Synchronized cells (1 × 10_7_) were harvested by trypsinization in the presence of appropriate compounds to maintain cell cycle phase and protein depletion. Cells were washed with PBS, fixed in 1% formaldehyde (Thermo Fisher Scientific, 28906) for 10 min at room temperature (RT) and quenched with 125 mM glycine for 5 min at RT. Pellets were collected and stored at -70°C. Unless otherwise stated, all centrifugation steps were performed at 1,100 x g for 1 min. Affi-prep Protein A beads (Bio-Rad, 1560005) were blocked overnight at 4°C in blocking buffer (20 mM Tris-HCl pH 8.0, 2 mM EDTA pH 8.0, 1% Triton-X, 150 mM NaCl, 1 mM phenylmethanesulfonylfluoride (PMSF; Sigma-Aldrich, 93482), 0.1 mg/ml BSA), washed and resuspended in dilution buffer (blocking buffer without BSA). Fixed cells were washed in ice-cold lysis buffer (50 mM Tris-HCl pH 8.0, 10 mM EDTA pH 8.0, 1% SDS, 1 mM PMSF (Sigma-Aldrich, 93482), 1-fold protease inhibitor cocktail (Roche, 11836170001)) and collected by centrifugation (1,700 x g for 2 min at 4°C). Chromatin was fragmented to an average size of 200-500 bp using Diagenode Bioruptor (high intensity, 12 cycles, 30 s on/30 s off). After removing a 10 µl aliquot as input control, chromatin was diluted 1:10 in dilution buffer and pre-cleared with blocked Protein A beads for 1h at 4°C. Pre-cleared chromatin was incubated overnight at 4°C with the following antibodies: 0.3 µg CTCF (Cell Signaling Technology, 2899), 10 µg RAD21 (Abcam, ab992), 10 µg Sororin (Abcam, ab192237). Chromatin-antibody complexes were captured with Protein A beads for 3 h at 4°C. Beads were washed twice with each of the following buffers: low-salt buffer (20 mM Tris-HCl pH 8.0, 2 mM EDTA pH 8.0, 1% Triton-X, 150 mM NaCl, 0.1% SDS, 1 mM PMSF (Sigma-Aldrich, 93482)), high salt buffer (20 mM Tris-HCl pH 8.0, 2 mM EDTA pH 8.0, 1 % Triton-X, 500 mM NaCl, 0.1% SDS, 1 mM PMSF (Sigma-Aldrich, 93482)), LiCl buffer (10 mM Tris-HCl pH 8.0, 2 mM EDTA pH 8.0, 250 mM LiCl, 0.5% NP-40, 0.5% sodium deoxycholate), and TE buffer (10 mM Tris-HCl pH 8.0, 1 mM EDTA pH 8.0). Chromatin was eluted (25 mM Tris-HCl pH 8.0, 5 mM EDTA pH 8.0, 0.5% SDS) at 65°C for 20 min with shaking (1,200 rpm). ChIP and input samples were treated with RNAse A (100 mg/ml, Qiagen, 19101) and Proteinase K (20 mg/ml, Qiagen, 19131) for 1 h at 37°C, followed by overnight incubation at 65°C to reverse crosslinks. DNA was purified using the Monarch PCR & DNA Cleanup kit (NEB, T1030). Sequencing libraries were prepared using NEBNext Ultra II DNA Library Prep Kit (NEB E7645) and sequenced on Illumina NextSeq 550 or NovaSeq X platforms. All antibodies used in this study are listed in Table S2.

### ChIP-seq data processing

Sequencing reads were processed using the Nextflow _108_ pipeline nf-core/chipseq v1.2.1 _109,110_. To ensure consistent read lengths across sequencing runs, reads were trimmed to 75 bp (--hard_trim5 75) and processed in single-end mode. Following read alignment to the human genome (hg19), replicate BAM files were merged using SAMtools v1.15 _111_. ChIP-seq signal tracks were generated in bigWig _112_ format using deepTools v3.5.4 _113_. For this step, mapped reads were extended to the fragment size estimated with MACS2 v2.2.9.1 _114_. Coverage was calculated in 50 bp genomic bins, excluding blacklisted regions _115_ and duplicate reads, followed by normalization to counts per million (CPM). Downstream analyses utilized signal tracks from immunoprecipitated (IP) samples only.

### ChIP-seq peak calling

ChIP-seq peaks were called using MACS2 with default parameters, utilizing immunoprecipitated (IP) samples as treatment and corresponding Input samples as control. The effective genome size was set to 2,736,124,898 (hg19), corresponding to unique mappers with 75 bp read length. To define robust peaks set, calls overlapping blacklisted regions were removed using BEDTools v2.31.1 _116_, and the remaining peaks were filtered by q-value threshold of < 1e-5. To ensure reproducibility, peaks were initially called on merged replicates; the final set retained only those merged peaks that overlapped with peaks called independently in at least two biological replicates.

### ChIP-seq average profiles

Average ChIP-seq profiles centered at regions of interest (TAD boundaries and CTCF peaks) were computed using pybbi v0.4.2 _117_, For each region, ChIP-seq signals were extracted from bigWig files within defined genomic windows and aggregated into a fixed number of equally sized bins; specific window sizes and resolutions are indicated in the corresponding figure legends. To compensate for regional copy-number variation (CNV), the signal in each bin was divided by the corresponding copy number estimated from Hi-C data (published scsHi-C “all”, 24hG2, 100 Kb resolution) using NeoLoopFinder v0.4.3 _118_. Bin-wise signals of individual regions were averaged and the genome-wide mean was subtracted from the averaged profiles to represent signal enrichment.

### ChIP-seq stackline profiles

To generate stackline profiles (heatmaps), ChIP-seq signal matrices were processed identically to the average profiles, including CNV correction and genome-wide mean subtraction. The rows of the resulting matrices were sorted randomly.

### Hi-C data processing

Sequencing reads were processed using the scsHi-C pipeline _119_ to generate .pairs _120_ files containing genomic contact coordinates. Replicate files were merged using pairtools v1.1.3 _121_. Following sister chromatid assignment, contacts were classified as cis sister, trans sister, or undefined, and separate .pairs files were generated for cis sister, trans sister, and all contacts. These files were converted into contact matrices at 1 kb resolution in cooler _122_ format using cooler v0.10.2 _123_. The cis sister and trans sister matrices were summed to generate a combined sister map. All resulting matrices were aggregated into multiple resolutions. The combined sister and all-contact maps were balanced by iterative correction _124_ with default parameters. Balancing weights derived from the combined sister maps were subsequently applied to the cis sister and trans sister maps.

To compare Hi-C map features across cell-cycle stages (Figure 1A–C), the G2-phase and G1-phase contact maps were downsampled to match the total number of contacts in the G2 ΔSororin map and the individual G2 biological replicates.

### TAD calling and comparison of annotations

The hierarchy of topologically associated domains was annotated using OnTAD v1.2 _43_ on scsHi-C maps of “all” contacts. To compare TAD architectures across cell cycle stages (Figure 1C), TADs were called on downsampled contact maps at 20Kb resolution using parameters -minsz 2, -maxsz 200, and -penalty 1e-1. TAD annotations were compared using the adjusted Rand Index (ARI) _42,43_. Briefly, each pixel in the contact map (up to 2 Mb genomic separation) was assigned the ID of the lowest-level TAD covering it. These pixel assignments were treated as clusters, and the similarity between clusterings from different maps was quantified using ARI.

To annotate hierarchical levels of TAD boundaries (Figure 2 and onwards), TADs were called on the full G2-phase contact map at 10 Kb resolution (-minsz 5, -maxsz 400, and -penalty 1e-1). To ensure robustness, the boundary list was filtered to exclude artifacts following the logic of the cooltools _125_ insulation method. Boundaries overlapping or adjacent to low-coverage bins were discarded. Additionally, boundaries were required to have at least 2/3 valid pixels in local insulation diamonds (+-50 kb, +-100 kb, and +-250 kb). Each valid boundary was assigned a hierarchical level corresponding to the highest level (smallest level number) of the TADs it demarcates. Boundaries were subsequently classified into three categories: high (level 1), medium (levels 2–3), and low (levels 4+).

### Average Hi-C contact maps

Average Hi-C contact maps (pileups) centered at TAD boundaries were calculated using cooltools v0.7.1. Briefly, contact submatrices centered on regions of interest were extracted and averaged pixel-wise. The first two diagonals were excluded to minimize Hi-C artifacts. To generate observed-over-expected pileups, the averaged observed contact frequencies were divided by the expected contact frequency (genome-wide average) for each genomic separation distance.

### Hi-C contact probability curves

Distance-dependent contact probability (*P(s)*) curves for cis- and trans-sister interactions were derived using pairtools. Contacts were aggregated into 72 geometrically spaced bins spanning genomic distances from 1 bp to 1 Gb. The contact frequency in each bin was normalized by the area of covered base pairs. To account for variation in sequencing depth, the cis- and trans-sister curves from each sample were scaled jointly, such that the total area under both curves summed to 1 across genomic separations of 10 kb to 200 Mb.

### Insulation score analysis

The insulation of cis- and trans-sister contacts was estimated using the insulation score, modified from _12_. Briefly, the score is calculated in sliding genomic windows as the negative log2 ratio of intra-window contacts spanning the window center to the total number of intra-window contacts. The resulting values are assigned to the window centers. Finally, to correct for differences in distance-decay effects (*P(s)*) between conditions, the genome-wide median is subtracted.

To profile the insulation landscape across cell-cycle stages (Figure 1B), insulation scores were computed from downsampled Hi-C maps of “all” contacts at 20kb resolution using a +-500Kb sliding window, excluding the first two diagonals. To profile insulation of sister-specific contacts in G2 phase, insulation scores were computed from full scsHi-C maps at 10kb resolution using a +-500Kb sliding window. For cis-sister insulation, the first two diagonals were excluded to minimize Hi-C artifacts. For trans-sister insulation, the first 15 diagonals were excluded to mitigate the influence of short-range trans-sister contact enrichment or depletion at boundaries. Average profiles of insulation scores at TAD boundaries of different levels (Figure 3 and onwards) were generated by aggregating genome-wide insulation score tracks in +-500kb windows centered on TAD boundaries.

### Assignment of CTCF sites to TAD boundaries

To assign CTCF sites to TAD boundaries, boundary coordinates were expanded to capture the extended transition zones characteristic of mammalian genomes_126_ and the associated clustering of CTCF sites _60_. Specifically, boundary regions were defined as 50 kb intervals centered on the original boundary calls (+-25 kb). This window size was selected as the maximum width that accommodates the extended boundary zone while ensuring that adjacent boundary regions remain non-overlapping. Reproducible CTCF peaks were then assigned to boundaries by intersecting peak coordinates with these expanded intervals.

### Statistical testing, data analysis, and visualization

Sequencing data were obtained from at least two biological replicates for each condition. Statistical testing was restricted to peak calling. In boxplots, the central line represents the median, the box limits indicate the first and third quartiles, and the whiskers extend to the 5th and 95th percentiles.

Analyses were performed in Python v3.10 using standard libraries (NumPy v1.25.2 _127_, Pandas v1.5.3 _128_, SciPy v1.15.2 _129_) and genomic analysis packages (bioframe v0.8.0 _130_, cooler, cooltools, pybbi, pybigtools v0.2.4 _117,122,123,125,131_. Data exploration and visualization were conducted using HiCognition _132_, IGV _133,134_, HiGlass _135_ via resgen.io, Matplotlib v3.9.4 _136_, and seaborn v0.13.2 _137_.

## Data and software availability

Raw sequencing data have been deposited in the ENA database under accession number <u>PRJEB108635</u>. Processed sequencing data have been deposited in the BioStudies database under accession number <u>S-BSST2754</u>. The code used for the analysis is available on GitHub at https://github.com/gerlichlab/takacs_mylarshchikov_et_al. To ensure reproducibility, a containerized JupyterLab _138_ environment including all software dependencies is available at https://github.com/gerlichlab/replchromconf-jupyterlab. Additionally, ChIP-seq track generation and peak calling were implemented as a Nextflow pipeline peakflow, which is available at https://github.com/gerlichlab/peakflow.

## Published datasets used

All published datasets used in this study are listed in Table S3.

---

**Figure S1.**
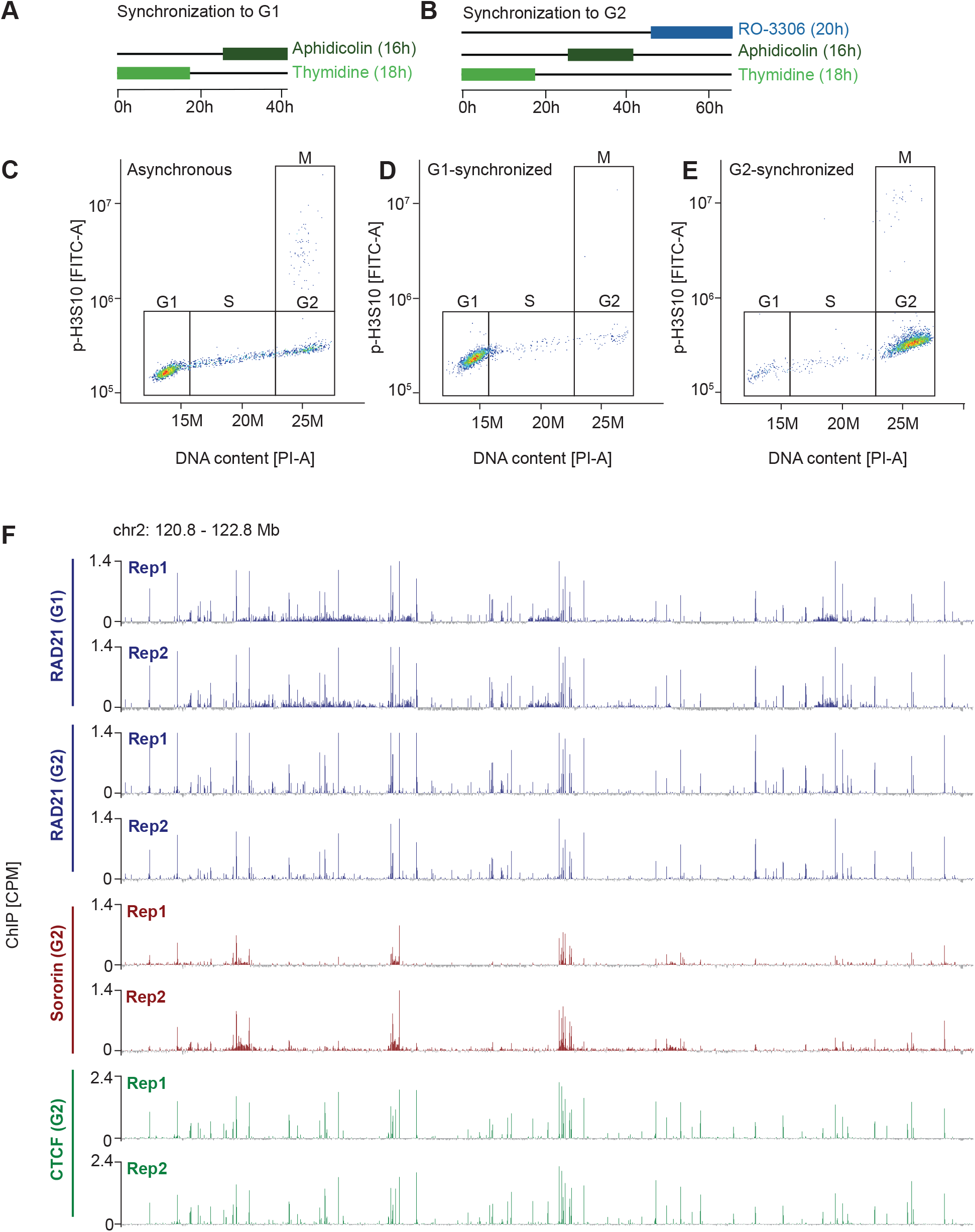
Characterization of cohesin distribution in G1 and G2 cell cycle phase. (**A-B**) Schematic of cell synchronization to G1 (A) and G2 (B) cell cycle phase. (**C-E**) Representative cell cycle profiles (phospho-H3S10 vs. DNA content) of asynchronous (C), G1-synchronized (D) and G2-synchronized (E) cells. (**F**) RAD21 ChIP-Seq signal tracks (1 kb resolution) from G1 cells and RAD21, Sororin and CTCF ChIP-Seq tracks from G2 cells at the locus shown in Figure 1A. Individual biological replicates are shown. RAD21 G1 and G2 replicates were downsampled to matching sequencing depth (60 mln). ChIP-Seq data are CPM-normalized and centered by subtracting the genome-wide mean.

**Figure S2.**
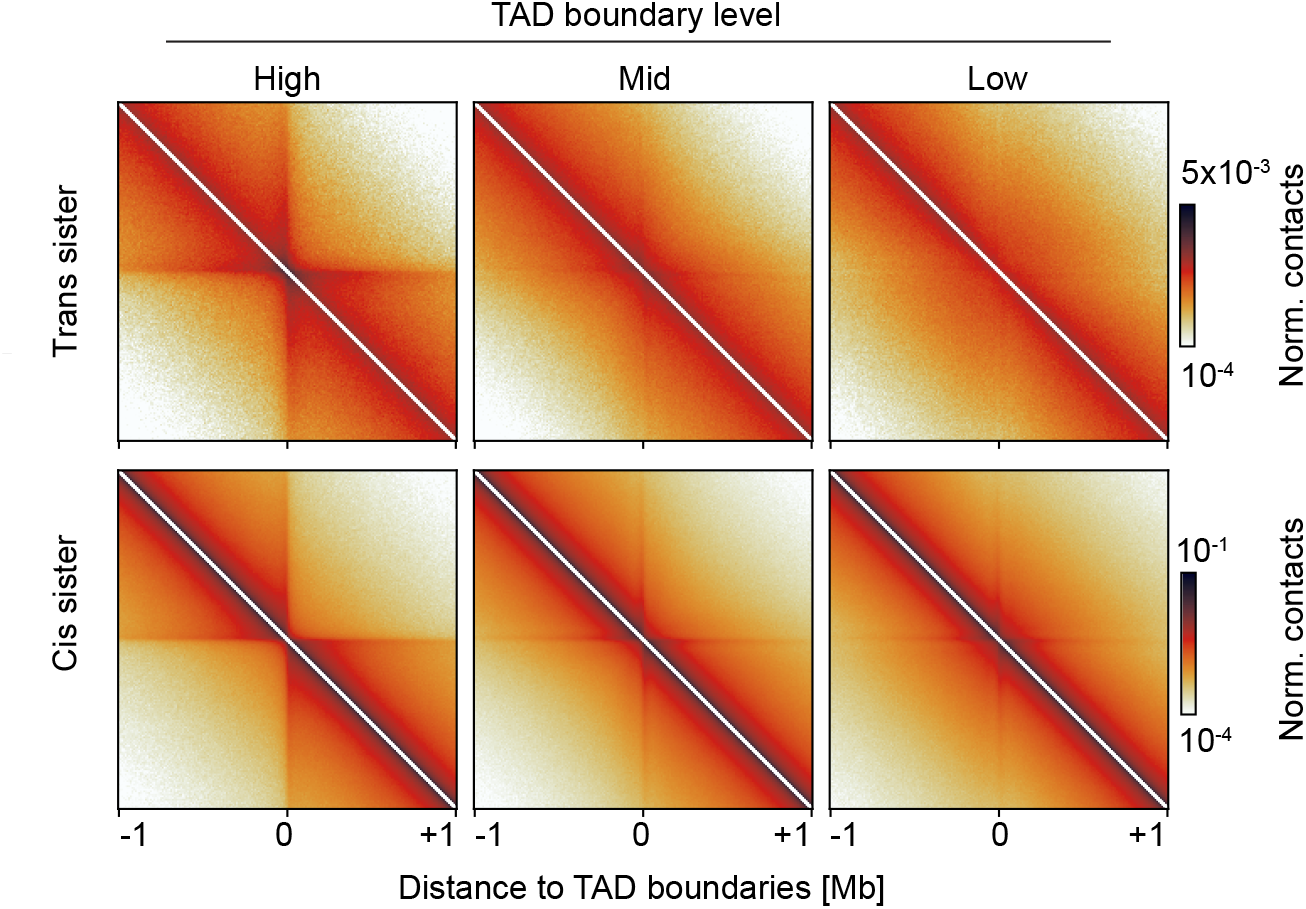
Trans sister contacts accumulate at high-level boundaries. Average maps of trans sister (upper row) and cis sister (lower row) scsHi-C contacts (10 kb resolution) at ±1 Mb regions centered at high-, mid-, and low-level boundaries. Wild type scsHi-C data reanalyzed from^33^.

**Figure S3.**
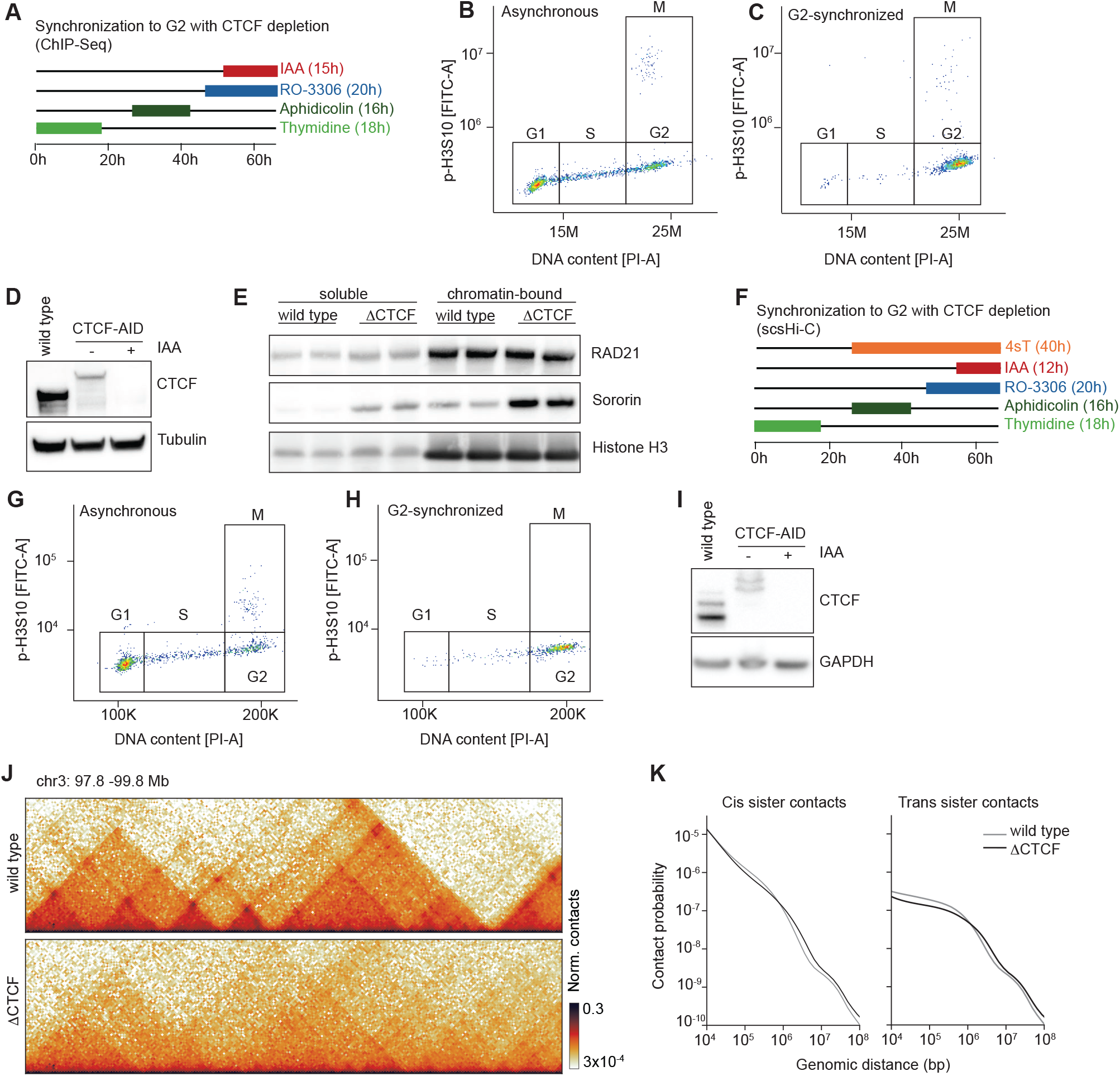
Characterization of CTCF depletion for scsHi-C and ChIP-Seq experiments. (**A**) Schematic of G2 cell synchronization and CTCF depletion for ChIP-Seq. (**B-C**) Representative cell cycle profiles (phospho-H3S10 vs. DNA content) of asynchronous (B) and G2-synchronized (C) cells corresponding to (A). (**D**) Western blot confirming CTCF depletion in G2 cells corresponding to (A-C). (**E)** Western blot confirming the presence of Sororin on chromatin in G2 ΔCTCF cells as shown in (A, C). Data are representative of n=2 independent experiments. (**F**) Schematic of G2 cell synchronization and CTCF depletion for scsHi-C. (**G-H**) Representative cell cycle profiles (phospho-H3S10 vs. DNA content) of asynchronous (G) and G2-synchronized (H) cells corresponding to (F). (**I**) Western blot confirming CTCF depletion in G2 cells corresponding to (F-H). (**J**) Representative Hi-C contact maps (10 kb resolution, 1 Mb separation) from wild type and CTCF-depleted (ΔCTCF) cells. (**K**) Distance-dependent contact probability (P(s)) curves for cis and trans sister scsHi-C-contacts from G2 WT and ΔCTCF cells. scsHi-C data for ΔCTCF are merged from n=5 independent experiments. Wild type scsHi-C data reanalyzed from^33^.

**Figure S4.**
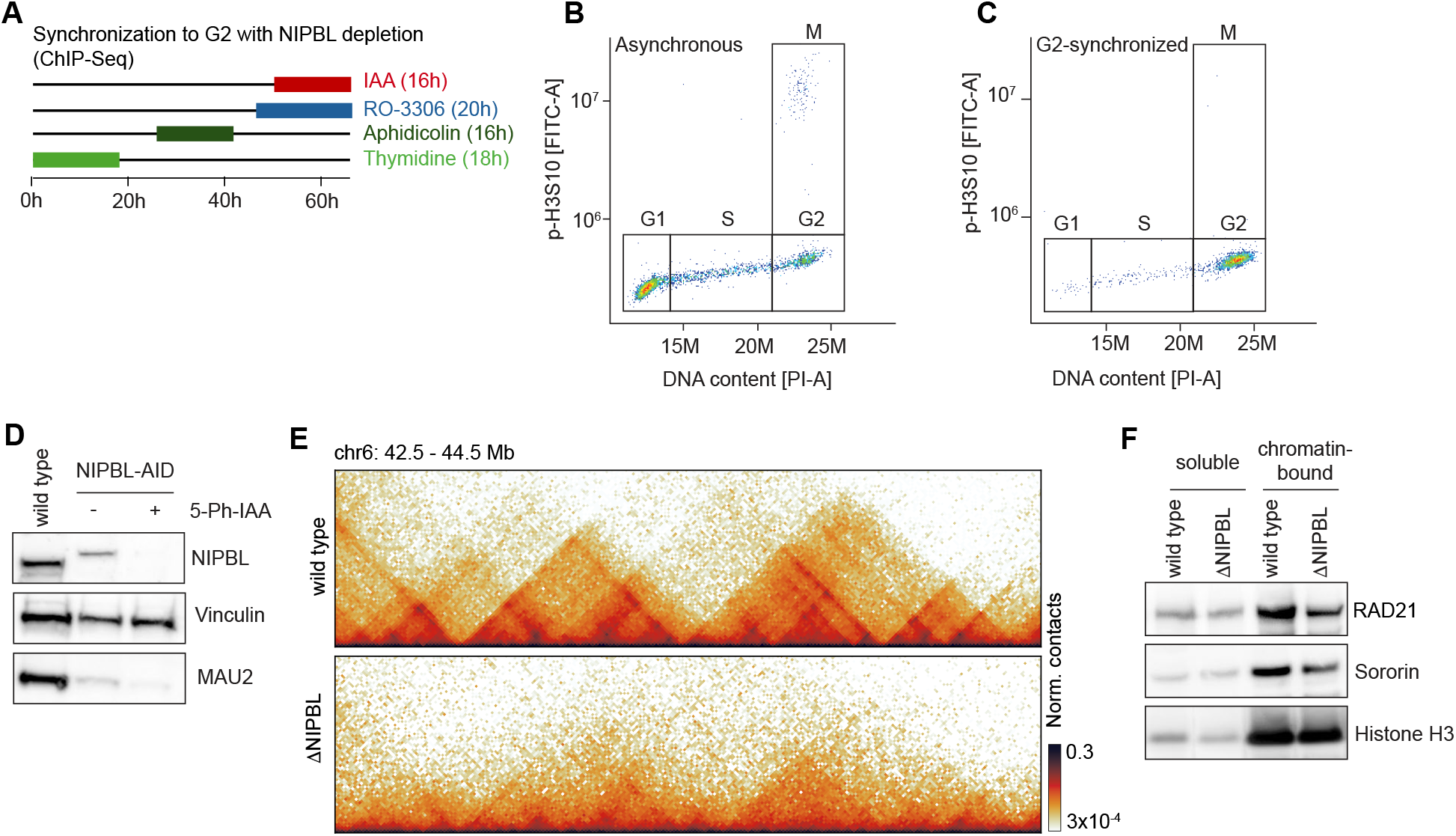
Characterization of NIPBL depletion for ChIP-Seq experiments. (**A**) Schematic of G2 synchronization and NIPBL depletion for ChIP-Seq. (**B-C**) Representative cell cycle profiles (phospho-H3S10 vs. DNA content) of asynchronous (B) and G2-synchronized (C) cells corresponding to (A). (**D**) Western blot confirming NIPBL depletion in G2 cells corresponding to (A-C). (**E**) Representative Hi-C contact maps (10 kb resolution, 1 Mb separation) from wild type and NIPBL-depleted (ΔNIPBL) cells. (**F**) Western blot confirming the presence of Sororin on chromatin in G2 ΔNIPBL cells as shown in (A, C). Data are representative of n=2 independent experiments. Wild type scsHi-C data reanalyzed from^33^. ΔNIPBL scsHi-C data reanalyzed from^34^.

**Figure S5.**
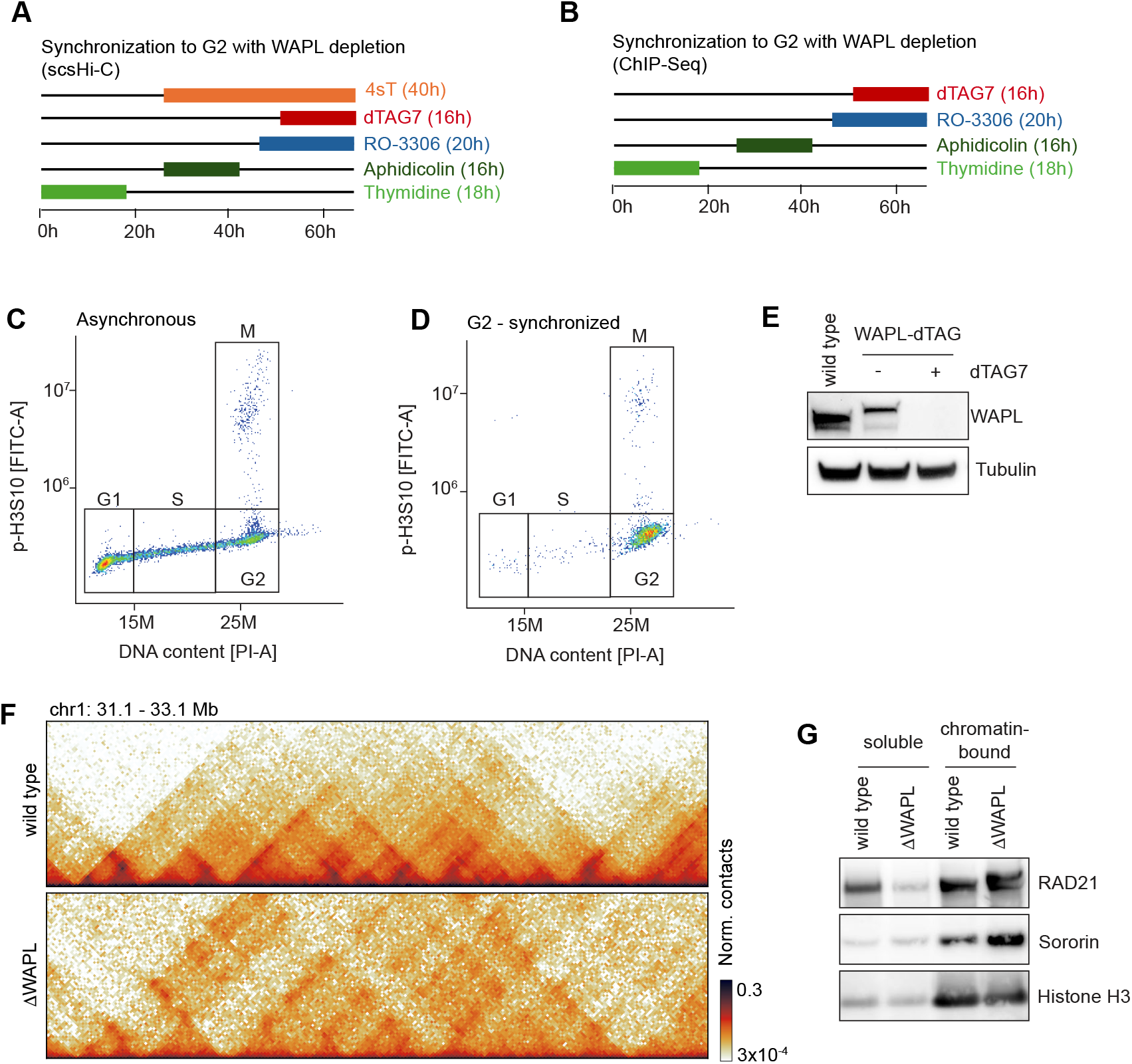
Characterization of WAPL depletion for scsHi-C and ChIP-Seq experiments. (**A-B**) Schematic of G2 synchronization and WAPL depletion for scsHi-C (A) and ChIP-Seq (B). (**C-D**) Representative cell cycle profiles (phospho-H3S10 vs. DNA content) of asynchronous (B) and G2-synchronized (C) cells corresponding to (A-B). (**E**) Western blot confirming WAPL depletion in G2 cells corresponding to (A, B, D). (**F**) Representative Hi-C contact maps (10 kb resolution, 1 Mb separation) from wild type and WAPL-depleted (ΔWAPL) cells. (**G**) Western blot confirming the presence of Sororin on chromatin in G2 ΔWAPL cells as shown in (B, D). scsHi-C data for ΔWAPL are merged from n=2 independent experiments. Wild type scsHi-C data reanalyzed from^33^.

**Figure S6.**
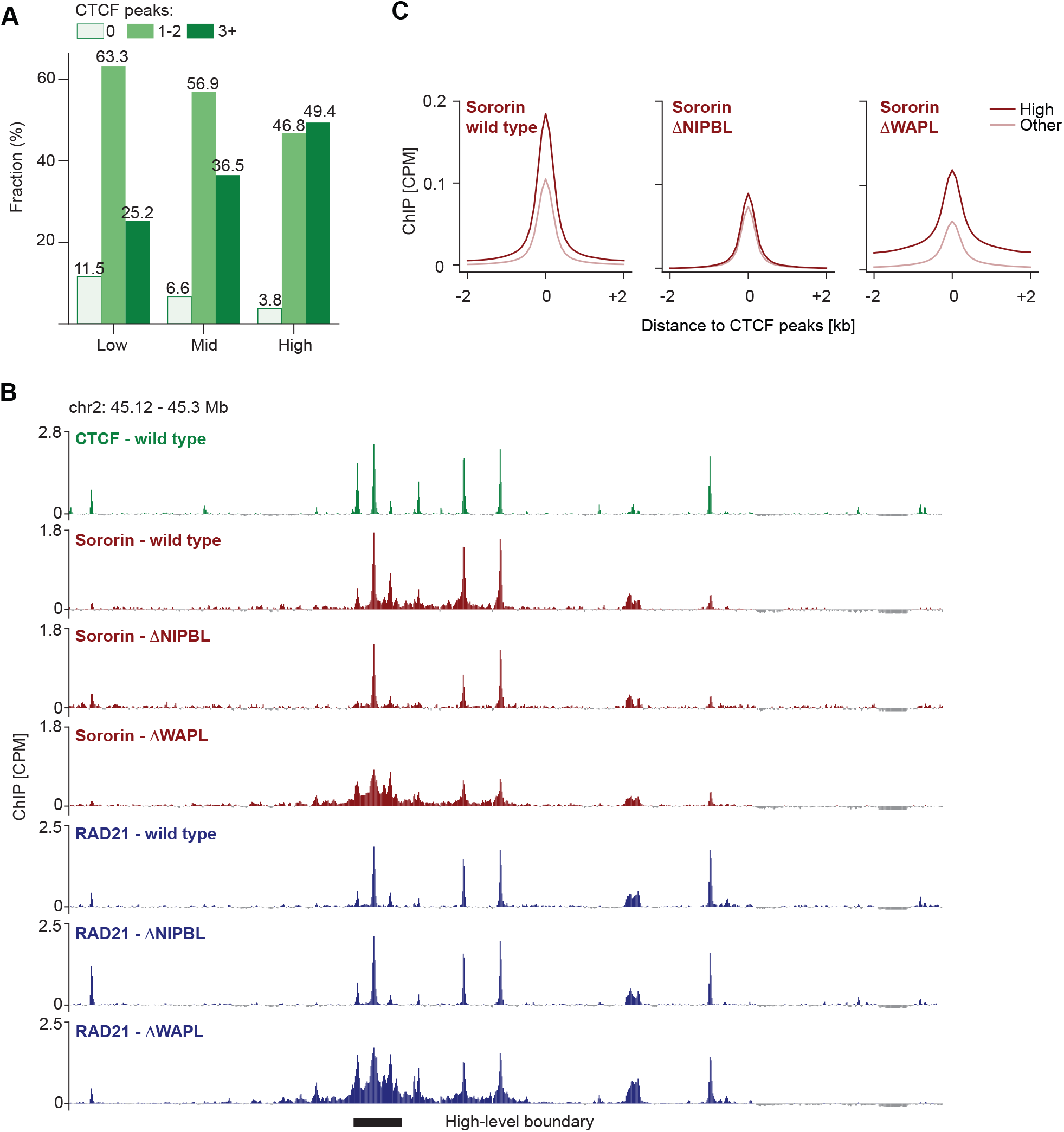
Characterization of CTCF clusters at high-level boundaries. (**A**) Fractions of low-, mid- and high-level boundaries containing 0, 1-2, or ≥3 CTCF peaks. (**B**) Representative ChIP-Seq tracks (200 bp resolution) of CTCF (green) from G2 wild type, and Sororin (red) and RAD21 (blue) from G2 wild type, NIPBL-depleted (ΔNIPBL) and WAPL-depleted (ΔWAPL) cells. Position of a high-level boundary (10 kb) is indicated below. (**C**) Average profile of Sororin ChIP-Seq signal (100 bp resolution) in G2 cells under wild type (left panel), ΔNIPBL (middle panel) and ΔWAPL (right panel) conditions at ±2 kb regions centered on isolated CTCF peaks (no additional CTCF peak within ±2.5kb) and stratified by location: high-level boundaries (High) (n=5989 peaks) or elsewhere in the genome (Other) (n=44600 peaks). ChIP-Seq data are CPM-normalized, centered by subtracting the genome-wide mean and represent merged data from n=3 (CTCF) or n=2 (Sororin, RAD21) independent experiments.

**Table S1.** A list of cell lines used in this study, and accompanying Gerlich lab IDs.

| Name | Genotype | Gerlich ID |
| --- | --- | --- |
| Wild type | HeLa Kyoto wild type | 0001 |
| NIPBL-AID | HeLa Kyoto, mEGFP-mAID-NIPBL OsTIR1(F74G)-SNAP-IRES-Blast (Batty et al. 2023) | 2045 |
| WAPL-dTAG | HeLa Kyoto, Blasticidin-P2A-2xHA-FKBP12F36V(dTAG)-WAPL (Batty et al. 2023) | 2096 |
| CTCF-AID | HeLa Kyoto, CTCF-mEGFP-AID (Wutz et al. 2017) | 1900 |
| CTCF-AID/SCC1-Halo | HeLa Kyoto, CTCF-AID-mKate2, Scc1-Halo (Wutz et al. 2025) | 1901 |

**Table S2.** A list of antibodies used in this study, including Gerlich lab ID, supplier, catalogue number and lot numbers. Abbreviations: CST – Cell Signaling Technology, SCBT – Santa Cruz Biotechnology.

| Antibody | Gerlich ID | Supplier | Catalogue # | Lot # |
| --- | --- | --- | --- | --- |
| Rabbit anti-CTCF | 350 | CST | 2899 | 3 |
| Rabbit anti-GAPDH | 229 | Abcam | ab9485 | GR3298316<br>GR3212164 |
| Rabbit anti-Histone H3 | 356 | CST | 4499 | 20 |
| Rabbit anti-SCC4/MAU2 | 441 | Abcam | ab183033 | GR3374490 |
| Rat anti-NIPBL | 334 | Absea | 010702F01 | 16060233915 |
| Rabbit anti-RAD21 (ChIP) | 385 | Abcam | ab992 | 1048469<br>GR3310168 |
| Rabbit anti-RAD21 (Immunoblotting) | 440 | Abcam | ab217678 | GR3362682 |
| Mouse anti-Sororin (Immunoblotting) | 432 | in-house |  | 3D5-G10 |
| Rabbit anti-Sororin (ChIP) | 371 | Abcam | ab192237 | 1065990<br>1025801<br>1004287<br>GR226228 |
| Rabbit anti-α-Tubulin | 395 | Abcam | ab52866 | GR3241238 |
| Rabbit anti-Vinculin | 330 | Abcam | ab129002 | GR221671 |
| Mouse anti-WAPL | 400 | SCBT | sc-365189 | H3120 |
| Goat anti-mouse-HRP | 294 | Bio-Rad | 1706516 | L005680 A |
| Goat anti-rabbit-HRP | 158 | Bio-Rad | 1706515 | 0000033407 |
| Goat anti-rat-HRP | 171 | Amersham | NA935 | 9559542 |

**Table S3.** A list of published datasets used in this study. Abbreviations: GEO – Gene Expression Omnibus.

| Dataset | GEO accession number | Publication |
| --- | --- | --- |
| HeLa Wild type G2, scsHi-C | GSE152373 | (Mitter et al. 2020) |
| HeLa Wild type prometaphase, scsHi-C | GSE152373 | (Mitter et al. 2020) |
| HeLa Sororin depleted G2, scsHi-C | GSE152373 | (Mitter et al. 2020) |
| HeLa NIPBL depleted G2, scsHi-C | GSE227694 | (Batty et al. 2023) |

## REFERENCES

1. Hoencamp, C., and Rowland, B.D. (2023). Genome control by SMC complexes. Nat. Rev. Mol. Cell Biol. 24, 633–650.

2. Ochs, F., and Gerlich, D.W. (2026). Organization of replicated chromosomes by DNA loops and sister chromatid cohesion. Nat. Rev. Mol. Cell Biol. 10.1038/s41580-025-00933-1.

3. Michaelis, C., Ciosk, R., and Nasmyth, K. (1997). Cohesins: chromosomal proteins that prevent premature separation of sister chromatids. Cell 91, 35–45.

4. Guacci, V., Koshland, D., and Strunnikov, A. (1997). A direct link between sister chromatid cohesion and chromosome condensation revealed through the analysis of MCD1 in S. cerevisiae. Cell 91, 47–57.

5. Davidson, I.F., Bauer, B., Goetz, D., Tang, W., Wutz, G., and Peters, J.-M. (2019). DNA loop extrusion by human cohesin. Science 366, 1338–1345.

6. Kim, Y., Shi, Z., Zhang, H., Finkelstein, I.J., and Yu, H. (2019). Human cohesin compacts DNA by loop extrusion. Science 366, 1345–1349.

7. Dixon, J.R., Selvaraj, S., Yue, F., Kim, A., Li, Y., Shen, Y., Hu, M., Liu, J.S., and Ren, B. (2012). Topological domains in mammalian genomes identified by analysis of chromatin interactions. Nature 485, 376–380.

8. Nora, E.P., Lajoie, B.R., Schulz, E.G., Giorgetti, L., Okamoto, I., Servant, N., Piolot, T., van Berkum, N.L., Meisig, J., Sedat, J., et al. (2012). Spatial partitioning of the regulatory landscape of the X-inactivation centre. Nature 485, 381–385.

9. Sexton, T., Yaffe, E., Kenigsberg, E., Bantignies, F., Leblanc, B., Hoichman, M., Parrinello, H., Tanay, A., and Cavalli, G. (2012). Three-dimensional folding and functional organization principles of the Drosophila genome. Cell 148, 458–472.

10. Rao, S.S.P., Huang, S.-C., Glenn St Hilaire, B., Engreitz, J.M., Perez, E.M., Kieffer-Kwon, K.-R., Sanborn, A.L., Johnstone, S.E., Bascom, G.D., Bochkov, I.D., et al. (2017). Cohesin Loss Eliminates All Loop Domains. Cell 171, 305–320.e24.

11. Schwarzer, W., Abdennur, N., Goloborodko, A., Pekowska, A., Fudenberg, G., Loe-Mie, Y., Fonseca, N.A., Huber, W., Haering, C.H., Mirny, L., et al. (2017). Two independent modes of chromatin organization revealed by cohesin removal. Nature 551, 51–56.

12. Wutz, G., Várnai, C., Nagasaka, K., Cisneros, D.A., Stocsits, R.R., Tang, W., Schoenfelder, S., Jessberger, G., Muhar, M., Hossain, M.J., et al. (2017). Topologically associating domains and chromatin loops depend on cohesin and are regulated by CTCF, WAPL, and PDS5 proteins. EMBO J. 36, 3573–3599.

13. Gassler, J., Brandão, H.B., Imakaev, M., Flyamer, I.M., Ladstätter, S., Bickmore, W.A., Peters, J.-M., Mirny, L.A., and Tachibana, K. (2017). A mechanism of cohesin-dependent loop extrusion organizes zygotic genome architecture. EMBO J. 36, 3600–3618.

14. Robson, M.I., Ringel, A.R., and Mundlos, S. (2019). Regulatory landscaping: How enhancer-promoter communication is sculpted in 3D. Mol. Cell 74, 1110–1122.

15. Karpinska, M.A., and Oudelaar, A.M. (2023). The role of loop extrusion in enhancer-mediated gene activation. Curr. Opin. Genet. Dev. 79, 102022.

16. Buckley, A., Vetralla, C., and Canzio, D. (2025). Gene clusters reveal fundamental principles of genome folding and transcriptional regulation. Annu. Rev. Cell Dev. Biol. 41, 579–603.

17. Gruber, S., Haering, C.H., and Nasmyth, K. (2003). Chromosomal cohesin forms a ring. Cell 112, 765–777.

18. Haering, C.H., Farcas, A.-M., Arumugam, P., Metson, J., and Nasmyth, K. (2008). The cohesin ring concatenates sister DNA molecules. Nature 454, 297–301.

19. Ochs, F., Green, C., Szczurek, A.T., Pytowski, L., Kolesnikova, S., Brown, J., Gerlich, D.W., Buckle, V., Schermelleh, L., and Nasmyth, K.A. (2024). Sister chromatid cohesion is mediated by individual cohesin complexes. Science 383, 1122–1130.

20. Sjögren, C., and Nasmyth, K. (2001). Sister chromatid cohesion is required for postreplicative double-strand break repair in Saccharomyces cerevisiae. Curr. Biol. 11, 991–995.

21. Sonoda, E., Matsusaka, T., Morrison, C., Vagnarelli, P., Hoshi, O., Ushiki, T., Nojima, K., Fukagawa, T., Waizenegger, I.C., Peters, J.M., et al. (2001). Scc1/Rad21/Mcd1 is required for sister chromatid cohesion and kinetochore function in vertebrate cells. Dev. Cell 1, 759–770.

22. Ström, L., Lindroos, H.B., Shirahige, K., and Sjögren, C. (2004). Postreplicative recruitment of cohesin to double-strand breaks is required for DNA repair. Mol. Cell 16, 1003–1015.

23. Teloni, F., Takacs, Z., Mitter, M., Langer, C.C.H., Prlesi, I., Steinacker, T.L., Reuter, V.P., Mylarshchikov, D., and Gerlich, D.W. (2025). Cohesin guides homology search during DNA repair using loops and sister chromatid linkages. Science 390, eadw0566.

24. Phipps, J., and Dubrana, K. (2022). DNA Repair in Space and Time: Safeguarding the Genome with the Cohesin Complex. Genes 13. 10.3390/genes13020198.

25. Tanaka, T., Fuchs, J., Loidl, J., and Nasmyth, K. (2000). Cohesin ensures bipolar attachment of microtubules to sister centromeres and resists their precocious separation. Nat. Cell Biol. 2, 492–499.

26. Oliveira, R.A., Hamilton, R.S., Pauli, A., Davis, I., and Nasmyth, K. (2010). Cohesin cleavage and Cdk inhibition trigger formation of daughter nuclei. Nat. Cell Biol. 12, 185–192.

27. Nasmyth, K., and Haering, C.H. (2009). Cohesin: its roles and mechanisms. Annu. Rev. Genet. 43, 525–558.

28. Yatskevich, S., Rhodes, J., and Nasmyth, K. (2019). Organization of Chromosomal DNA by SMC Complexes. Annu. Rev. Genet. 53, 445–482.

29. Davidson, I.F., and Peters, J.-M. (2021). Genome folding through loop extrusion by SMC complexes. Nat. Rev. Mol. Cell Biol. 10.1038/s41580-021-00349-7.

30. Corsi, F., Rusch, E., and Goloborodko, A. (2023). Loop extrusion rules: the next generation. Curr. Opin. Genet. Dev. 81, 102061.

31. Dekker, J., and Mirny, L.A. (2024). The chromosome folding problem and how cells solve it. Cell 187, 6424–6450.

32. Goloborodko, A., Imakaev, M.V., Marko, J.F., and Mirny, L. (2016). Compaction and segregation of sister chromatids via active loop extrusion. Elife 5. 10.7554/eLife.14864.

33. Mitter, M., Gasser, C., Takacs, Z., Langer, C.C.H., Tang, W., Jessberger, G., Beales, C.T., Neuner, E., Ameres, S.L., Peters, J.-M., et al. (2020). Conformation of sister chromatids in the replicated human genome. Nature 586, 139–144.

34. Batty, P., Langer, C.C., Takács, Z., Tang, W., Blaukopf, C., Peters, J.-M., and Gerlich, D.W. (2023). Cohesin-mediated DNA loop extrusion resolves sister chromatids in G2 phase. EMBO J. 42, e113475.

35. Chatzidaki, E.E., Powell, S., Dequeker, B.J.H., Gassler, J., Silva, M.C.C., and Tachibana, K. (2021). Ovulation suppression protects against chromosomal abnormalities in mouse eggs at advanced maternal age. Curr. Biol. 31, 4038–4051.e7.

36. Bastié, N., Chapard, C., Cournac, A., Nejmi, S., Mboumba, H., Gadal, O., Thierry, A., Beckouët, F., and Koszul, R. (2024). Sister chromatid cohesion halts DNA loop expansion. Mol. Cell 84, 1139–1148.e5.

37. Zhao, H., Shu, L., Qin, S., Lyu, F., Liu, F., Lin, E., Xia, S., Wang, B., Wang, M., Shan, F., et al. (2025). Extensive mutual influences of SMC complexes shape 3D genome folding. Nature 640, 543–553.

38. Samejima, K., Gibcus, J.H., Abraham, S., Cisneros-Soberanis, F., Samejima, I., Beckett, A.J., Pučeková, N., Abad, M.A., Spanos, C., Medina-Pritchard, B., et al. (2025). Rules of engagement for condensins and cohesins guide mitotic chromosome formation. Science 388, eadq1709.

39. Naumova, N., Imakaev, M., Fudenberg, G., Zhan, Y., Lajoie, B.R., Mirny, L.A., and Dekker, J. (2013). Organization of the mitotic chromosome. Science 342, 948–953.

40. Gibcus, J.H., Samejima, K., Goloborodko, A., Samejima, I., Naumova, N., Nuebler, J., Kanemaki, M.T., Xie, L., Paulson, J.R., Earnshaw, W.C., et al. (2018). A pathway for mitotic chromosome formation. Science 359. 10.1126/science.aao6135.

41. Nagano, T., Lubling, Y., Várnai, C., Dudley, C., Leung, W., Baran, Y., Mendelson Cohen, N., Wingett, S., Fraser, P., and Tanay, A. (2017). Cell-cycle dynamics of chromosomal organization at single-cell resolution. Nature 547, 61–67.

42. Hubert, L., and Arabie, P. (1985). Comparing partitions. J. Classif. 2, 193–218.

43. An, L., Yang, T., Yang, J., Nuebler, J., Xiang, G., Hardison, R.C., Li, Q., and Zhang, Y. (2019). OnTAD: hierarchical domain structure reveals the divergence of activity among TADs and boundaries. Preprint, 10.1186/s13059-019-1893-y https://doi.org/10.1186/s13059-019-1893-y.

44. Rankin, S., Ayad, N.G., and Kirschner, M.W. (2005). Sororin, a substrate of the anaphase-promoting complex, is required for sister chromatid cohesion in vertebrates. Mol. Cell 18, 185–200.

45. Schmitz, J., Watrin, E., Lénárt, P., Mechtler, K., and Peters, J.-M. (2007). Sororin is required for stable binding of cohesin to chromatin and for sister chromatid cohesion in interphase. Curr. Biol. 17, 630–636.

46. Ladurner, R., Kreidl, E., Ivanov, M.P., Ekker, H., Idarraga-Amado, M.H., Busslinger, G.A., Wutz, G., Cisneros, D.A., and Peters, J.-M. (2016). Sororin actively maintains sister chromatid cohesion. EMBO J. 35, 635–653.

47. Gerlich, D., Koch, B., Dupeux, F., Peters, J.-M., and Ellenberg, J. (2006). Live-cell imaging reveals a stable cohesin-chromatin interaction after but not before DNA replication. Curr. Biol. 16, 1571–1578.

48. Holzmann, J., Politi, A.Z., Nagasaka, K., Hantsche-Grininger, M., Walther, N., Koch, B., Fuchs, J., Dürnberger, G., Tang, W., Ladurner, R., et al. (2019). Absolute quantification of cohesin, CTCF and their regulators in human cells. Elife 8. 10.7554/eLife.46269.

49. Peters, J.-M., and Nishiyama, T. (2012). Sister chromatid cohesion. Cold Spring Harb. Perspect. Biol. 4. 10.1101/cshperspect.a011130.

50. Wendt, K.S., Yoshida, K., Itoh, T., Bando, M., Koch, B., Schirghuber, E., Tsutsumi, S., Nagae, G., Ishihara, K., Mishiro, T., et al. (2008). Cohesin mediates transcriptional insulation by CCCTC-binding factor. Nature 451, 796–801.

51. Parelho, V., Hadjur, S., Spivakov, M., Leleu, M., Sauer, S., Gregson, H.C., Jarmuz, A., Canzonetta, C., Webster, Z., Nesterova, T., et al. (2008). Cohesins functionally associate with CTCF on mammalian chromosome arms. Cell 132, 422–433.

52. Busslinger, G.A., Stocsits, R.R., van der Lelij, P., Axelsson, E., Tedeschi, A., Galjart, N., and Peters, J.-M. (2017). Cohesin is positioned in mammalian genomes by transcription, CTCF and Wapl. Nature 544, 503–507.

53. Nora, E.P., Goloborodko, A., Valton, A.-L., Gibcus, J.H., Uebersohn, A., Abdennur, N., Dekker, J., Mirny, L.A., and Bruneau, B.G. (2017). Targeted degradation of CTCF decouples local insulation of chromosome domains from genomic compartmentalization. Cell 169, 930–944.e22.

54. Mitter, M., Takacs, Z., Köcher, T., Micura, R., Langer, C.C.H., and Gerlich, D.W. (2022). Sister chromatid-sensitive Hi-C to map the conformation of replicated genomes. Nat. Protoc. 17, 1486–1517.

55. Rao, S.S.P., Huntley, M.H., Durand, N.C., Stamenova, E.K., Bochkov, I.D., Robinson, J.T., Sanborn, A.L., Machol, I., Omer, A.D., Lander, E.S., et al. (2014). A 3D map of the human genome at kilobase resolution reveals principles of chromatin looping. Cell 159, 1665–1680.

56. Fudenberg, G., Imakaev, M., Lu, C., Goloborodko, A., Abdennur, N., and Mirny, L.A. (2016). Formation of chromosomal domains by loop extrusion. Cell Rep. 15, 2038–2049.

57. Sanborn, A.L., Rao, S.S.P., Huang, S.-C., Durand, N.C., Huntley, M.H., Jewett, A.I., Bochkov, I.D., Chinnappan, D., Cutkosky, A., Li, J., et al. (2015). Chromatin extrusion explains key features of loop and domain formation in wild-type and engineered genomes. Proc. Natl. Acad. Sci. U. S. A. 112, E6456–65.

58. Nuebler, J., Fudenberg, G., Imakaev, M., Abdennur, N., and Mirny, L.A. (2018). Chromatin organization by an interplay of loop extrusion and compartmental segregation. Proc. Natl. Acad. Sci. U. S. A. 115, E6697–E6706.

59. Davidson, I.F., Barth, R., Zaczek, M., van der Torre, J., Tang, W., Nagasaka, K., Janissen, R., Kerssemakers, J., Wutz, G., Dekker, C., et al. (2023). CTCF is a DNA-tension-dependent barrier to cohesin-mediated loop extrusion. Nature 616, 822–827.

60. Chang, L.-H., Ghosh, S., Papale, A., Luppino, J.M., Miranda, M., Piras, V., Degrouard, J., Edouard, J., Poncelet, M., Lecouvreur, N., et al. (2023). Multi-feature clustering of CTCF binding creates robustness for loop extrusion blocking and Topologically Associating Domain boundaries. Nat. Commun. 14, 5615.

61. Zhang, H., Shi, Z., Banigan, E.J., Kim, Y., Yu, H., Bai, X.-C., and Finkelstein, I.J. (2023). CTCF and R-loops are boundaries of cohesin-mediated DNA looping. Mol. Cell 83, 2856–2871.e8.

62. Li, Y., Haarhuis, J.H.I., Sedeño Cacciatore, Á., Oldenkamp, R., van Ruiten, M.S., Willems, L., Teunissen, H., Muir, K.W., de Wit, E., Rowland, B.D., et al. (2020). The structural basis for cohesin-CTCF-anchored loops. Nature 578, 472–476.

63. Nora, E.P., Caccianini, L., Fudenberg, G., So, K., Kameswaran, V., Nagle, A., Uebersohn, A., Hajj, B., Saux, A.L., Coulon, A., et al. (2020). Molecular basis of CTCF binding polarity in genome folding. Nat. Commun. 11, 5612.

64. Panizza, S., Tanaka, T., Hochwagen, A., Eisenhaber, F., and Nasmyth, K. (2000). Pds5 cooperates with cohesin in maintaining sister chromatid cohesion. Curr. Biol. 10, 1557–1564.

65. Sumara, I., Vorlaufer, E., Gieffers, C., Peters, B.H., and Peters, J.M. (2000). Characterization of vertebrate cohesin complexes and their regulation in prophase. J. Cell Biol. 151, 749–762.

66. Hartman, T., Stead, K., Koshland, D., and Guacci, V. (2000). Pds5p is an essential chromosomal protein required for both sister chromatid cohesion and condensation in Saccharomyces cerevisiae. J. Cell Biol. 151, 613–626.

67. Kikuchi, S., Borek, D.M., Otwinowski, Z., Tomchick, D.R., and Yu, H. (2016). Crystal structure of the cohesin loader Scc2 and insight into cohesinopathy. Proc. Natl. Acad. Sci. U. S. A. 113, 12444–12449.

68. Petela, N.J., Gligoris, T.G., Metson, J., Lee, B.-G., Voulgaris, M., Hu, B., Kikuchi, S., Chapard, C., Chen, W., Rajendra, E., et al. (2018). Scc2 is a potent activator of cohesin’s ATPase that promotes loading by binding Scc1 without Pds5. Mol. Cell 70, 1134–1148.e7.

69. Ciosk, R., Shirayama, M., Shevchenko, A., Tanaka, T., Toth, A., Shevchenko, A., and Nasmyth, K. (2000). Cohesin’s binding to chromosomes depends on a separate complex consisting of Scc2 and Scc4 proteins. Mol. Cell 5, 243–254.

70. Murayama, Y., and Uhlmann, F. (2014). Biochemical reconstitution of topological DNA binding by the cohesin ring. Nature 505, 367–371.

71. Wutz, G., Davidson, I.F., Banigan, E.J., Kawasumi, R., Stocsits, R.R., Tang, W., Nagasaka, K., Costantino, L., Jansen, R., Hirota, K., et al. (2025). PDS5 proteins control genome architecture by limiting the lifetime of cohesin-NIPBL complexes. Molecular Biology.

72. Shah, R., Tortora, M.M.C., Louafi, N., Rahmaninejad, H., Hansen, K.L., Anderson, E.C., Wen, D., Giorgetti, L., Fudenberg, G., and Nora, E.P. (2025). Dosage sensitivity of the loop extrusion rate confers tunability to genome folding while creating vulnerability to genetic disruption. Molecular Biology.

73. Srinivasan, M., Petela, N.J., Scheinost, J.C., Collier, J., Voulgaris, M., B Roig, M., Beckouët, F., Hu, B., and Nasmyth, K.A. (2019). Scc2 counteracts a Wapl-independent mechanism that releases cohesin from chromosomes during G1. Elife 8. 10.7554/eLife.44736.

74. Nasmyth, K.A., Lee, B.-G., Roig, M.B., and Löwe, J. (2023). What AlphaFold tells us about cohesin’s retention on and release from chromosomes. Elife 12. 10.7554/eLife.88656.

75. Kueng, S., Hegemann, B., Peters, B.H., Lipp, J.J., Schleiffer, A., Mechtler, K., and Peters, J.-M. (2006). Wapl controls the dynamic association of cohesin with chromatin. Cell 127, 955–967.

76. Gause, M., Misulovin, Z., Bilyeu, A., and Dorsett, D. (2010). Dosage-sensitive regulation of cohesin chromosome binding and dynamics by Nipped-B, Pds5, and Wapl. Mol. Cell. Biol. 30, 4940–4951.

77. Tedeschi, A., Wutz, G., Huet, S., Jaritz, M., Wuensche, A., Schirghuber, E., Davidson, I.F., Tang, W., Cisneros, D.A., Bhaskara, V., et al. (2013). Wapl is an essential regulator of chromatin structure and chromosome segregation. Nature 501, 564–568.

78. Huis in ’t Veld, P.J., Herzog, F., Ladurner, R., Davidson, I.F., Piric, S., Kreidl, E., Bhaskara, V., Aebersold, R., and Peters, J.-M. (2014). Characterization of a DNA exit gate in the human cohesin ring. Science 346, 968–972.

79. Chan, B., and Rubinstein, M. (2023). Theory of chromatin organization maintained by active loop extrusion. Proc. Natl. Acad. Sci. U. S. A. 120, e2222078120.

80. Kiefer, L., Chiosso, A., Langen, J., Buckley, A., Gaudin, S., Rajkumar, S.M., Servito, G.I.F., Cha, E.S., Vijay, A., Yeung, A., et al. (2023). WAPL functions as a rheostat of Protocadherin isoform diversity that controls neural wiring. Science 380, eadf8440.

81. Haarhuis, J.H.I., van der Weide, R.H., Blomen, V.A., Yáñez-Cuna, J.O., Amendola, M., van Ruiten, M.S., Krijger, P.H.L., Teunissen, H., Medema, R.H., van Steensel, B., et al. (2017). The cohesin release factor WAPL restricts chromatin loop extension. Cell 169, 693–707.e14.

82. Luppino, J.M., Park, D.S., Nguyen, S.C., Lan, Y., Xu, Z., Yunker, R., and Joyce, E.F. (2020). Cohesin promotes stochastic domain intermingling to ensure proper regulation of boundary-proximal genes. Nat. Genet. 52, 840–848.

83. Kentepozidou, E., Aitken, S.J., Feig, C., Stefflova, K., Ibarra-Soria, X., Odom, D.T., Roller, M., and Flicek, P. (2020). Clustered CTCF binding is an evolutionary mechanism to maintain topologically associating domains. Genome Biol. 21, 5.

84. Anania, C., Acemel, R.D., Jedamzick, J., Bolondi, A., Cova, G., Brieske, N., Kühn, R., Wittler, L., Real, F.M., and Lupiáñez, D.G. (2022). In vivo dissection of a clustered-CTCF domain boundary reveals developmental principles of regulatory insulation. Nat. Genet. 54, 1026–1036.

85. Rudnizky, S., Murray, P.J., Sørensen, E.W., Koenig, T.J.R., Pangeni, S., Merino-Urteaga, R., Chhabra, H., Caccianini, L., Davidson, I.F., Osorio-Valeriano, M., et al. (2025). Ultrafast CTCF dynamics control cohesin barrier function. Biophysics.

86. Chang, L.-H., and Noordermeer, D. (2024). Permeable TAD boundaries and their impact on genome-associated functions. Bioessays 46, e2400137.

87. Yeh, C.D., van de Venn, L., Kreutzer, S., Zheng, X., Cantos, N.C., Schröder, M., Hofmann, R., Gerbaldo, F.E., Clemens, A., Wienert, B., et al. (2025). Proximity determines donor candidacy during DNA double-stranded break homology directed repair. bioRxiv. 10.1101/2025.02.10.637161.

88. Marin-Gonzalez, A., Rybczynski, A.T., Nilavar, N.M., Nguyen, D., Li, A.G., Karwacki-Neisius, V., Zou, R.S., Avilés-Vázquez, F.J., Kanemaki, M.T., Scully, R., et al. (2025). Cohesin drives chromatin scanning during the RAD51-mediated homology search. Science 390, eadw1928.

89. Renkawitz, J., Lademann, C.A., and Jentsch, S. (2014). Mechanisms and principles of homology search during recombination. Nat. Rev. Mol. Cell Biol. 15, 369–383.

90. Haber, J.E. (2018). DNA repair: The search for homology. Bioessays 40, 1700229.

91. Arnould, C., Rocher, V., Finoux, A.-L., Clouaire, T., Li, K., Zhou, F., Caron, P., Mangeot, P.E., Ricci, E.P., Mourad, R., et al. (2021). Loop extrusion as a mechanism for formation of DNA damage repair foci. Nature 590, 660–665.

92. Piazza, A., Bordelet, H., Dumont, A., Thierry, A., Savocco, J., Girard, F., and Koszul, R. (2021). Cohesin regulates homology search during recombinational DNA repair. Nat. Cell Biol. 23, 1176–1186.

93. Kim, E., Barth, R., and Dekker, C. (2023). Looping the genome with SMC complexes. Annu. Rev. Biochem. 92, 15–41.

94. Dequeker, B.J.H., Scherr, M.J., Brandão, H.B., Gassler, J., Powell, S., Gaspar, I., Flyamer, I.M., Lalic, A., Tang, W., Stocsits, R., et al. (2022). MCM complexes are barriers that restrict cohesin-mediated loop extrusion. Nature 606, 197–203.

95. Borrie, M.S., Kraycer, P.M., and Gartenberg, M.R. (2023). Transcription-driven translocation of cohesive and non-cohesive cohesin in vivo. Mol. Cell. Biol. 43, 254–268.

96. Borrie, M.S., Campor, J.S., Joshi, H., and Gartenberg, M.R. (2017). Binding, sliding, and function of cohesin during transcriptional activation. Proc. Natl. Acad. Sci. U. S. A. 114, E1062–E1071.

97. Lengronne, A., Katou, Y., Mori, S., Yokobayashi, S., Kelly, G.P., Itoh, T., Watanabe, Y., Shirahige, K., and Uhlmann, F. (2004). Cohesin relocation from sites of chromosomal loading to places of convergent transcription. Nature 430, 573–578.

98. Banigan, E.J., Tang, W., van den Berg, A.A., Stocsits, R.R., Wutz, G., Brandão, H.B., Busslinger, G.A., Peters, J.-M., and Mirny, L.A. (2023). Transcription shapes 3D chromatin organization by interacting with loop extrusion. Proc. Natl. Acad. Sci. U. S. A. 120, e2210480120.

99. Glynn, E.F., Megee, P.C., Yu, H.-G., Mistrot, C., Unal, E., Koshland, D.E., DeRisi, J.L., and Gerton, J.L. (2004). Genome-wide mapping of the cohesin complex in the yeast Saccharomyces cerevisiae. PLoS Biol. 2, E259.

100. Ocampo-Hafalla, M., Muñoz, S., Samora, C.P., and Uhlmann, F. (2016). Evidence for cohesin sliding along budding yeast chromosomes. Open Biol. 6. 10.1098/rsob.150178.

101. Gerlich, D., Hirota, T., Koch, B., Peters, J.-M., and Ellenberg, J. (2006). Condensin I stabilizes chromosomes mechanically through a dynamic interaction in live cells. Curr. Biol. 16, 333–344.

102. Walther, N., Hossain, M.J., Politi, A.Z., Koch, B., Kueblbeck, M., Ødegård-Fougner, Ø., Lampe, M., and Ellenberg, J. (2018). A quantitative map of human Condensins provides new insights into mitotic chromosome architecture. J. Cell Biol. 217, 2309–2328.

103. Dey, A., Shi, G., Takaki, R., and Thirumalai, D. (2023). Structural changes in chromosomes driven by multiple condensin motors during mitosis. Cell Rep. 42, 112348.

104. Oomen, M.E., Hans A.S., Liu, Y., Darzacq, X., and Dekker, J. (2019). CTCF sites display cell cycle-dependent dynamics in factor binding and nucleosome positioning. Genome Res. 29, 236–249.

105. Hill, L., Ebert, A., Jaritz, M., Wutz, G., Nagasaka, K., Tagoh, H., Kostanova-Poliakova, D., Schindler, K., Sun, Q., Bönelt, P., et al. (2020). Wapl repression by Pax5 promotes V gene recombination by Igh loop extrusion. Nature 584, 142–147.

106. Peters, J.-M. (2021). How DNA loop extrusion mediated by cohesin enables V(D)J recombination. Curr. Opin. Cell Biol. 70, 75–83.

107. Zhang, Y., Zhang, X., Dai, H.-Q., Hu, H., and Alt, F.W. (2022). The role of chromatin loop extrusion in antibody diversification. Nat. Rev. Immunol. 22, 550–566.

## METHODS REFERENCES

108. Di Tommaso, P., Chatzou, M., Floden, E.W., Barja, P.P., Palumbo, E., and Notredame, C. (2017). Nextflow enables reproducible computational workflows. Nat. Biotechnol. 35, 316–319.

109. Ewels, P.A., Peltzer, A., Fillinger, S., Patel, H., Alneberg, J., Wilm, A., Garcia, M.U., Di Tommaso, P., and Nahnsen, S. (2020). The nf-core framework for community-curated bioinformatics pipelines. Nat. Biotechnol. 38, 276–278.

110. Patel, H., Wang, C., Ewels, P., Peltzer, A., Silva, T.C., Behrens, D., Garcia, M., mashehu, Rotholandus, Haglund, S., et al. (2020). nf-core/chipseq: nf-core/chipseq v1.2.1 - Platinum Mole (Zenodo) 10.5281/ZENODO.3966161.

111. Li, H., Handsaker, B., Wysoker, A., Fennell, T., Ruan, J., Homer, N., Marth, G., Abecasis, G., Durbin, R., and 1000 Genome Project Data Processing Subgroup (2009). The Sequence Alignment/Map format and SAMtools. Bioinformatics 25, 2078–2079.

112. Kent, W.J., Zweig, A.S., Barber, G., Hinrichs, A.S., and Karolchik, D. (2010). BigWig and BigBed: enabling browsing of large distributed datasets. Bioinformatics 26, 2204–2207.

113. Ramírez, F., Ryan, D.P., Grüning, B., Bhardwaj, V., Kilpert, F., Richter, A.S., Heyne, S., Dündar, F., and Manke, T. (2016). deepTools2: a next generation web server for deep-sequencing data analysis. Nucleic Acids Res. 44, W160–5.

114. Zhang, Y., Liu, T., Meyer, C.A., Eeckhoute, J., Johnson, D.S., Bernstein, B.E., Nusbaum, C., Myers, R.M., Brown, M., Li, W., et al. (2008). Model-based analysis of ChIP-Seq (MACS). Genome Biol. 9, R137.

115. Amemiya, H.M., Kundaje, A., and Boyle, A.P. (2019). The ENCODE blacklist: Identification of problematic regions of the genome. Sci. Rep. 9, 9354.

116. Quinlan, A.R., and Hall, I.M. (2010). BEDTools: a flexible suite of utilities for comparing genomic features. Bioinformatics 26, 841–842.

117. Abdennur, N. (2025). pybbi (Zenodo) 10.5281/ZENODO.17485681.

118. Wang, X., Xu, J., Zhang, B., Hou, Y., Song, F., Lyu, H., and Yue, F. (2021). Genome-wide detection of enhancer-hijacking events from chromatin interaction data in rearranged genomes. Nat. Methods 18, 661–668.

119. Langer, C.C.H. (2021). scsHi-C preprocessing Nextflow pipeline 10.5281/zenodo.5742764.

120. Lee, S., Bakker, C.R., Vitzthum, C., Alver, B.H., and Park, P.J. (2022). Pairs and Pairix: a file format and a tool for efficient storage and retrieval for Hi-C read pairs. Bioinformatics 38, 1729–1731.

121. Open2C, Abdennur, N., Fudenberg, G., Flyamer, I.M., Galitsyna, A.A., Goloborodko, A., Imakaev, M., and Venev, S.V. (2023). Pairtools: from sequencing data to chromosome contacts. bioRxivorg. 10.1101/2023.02.13.528389.

122. Abdennur, N., and Mirny, L.A. (2020). Cooler: scalable storage for Hi-C data and other genomically labeled arrays. Bioinformatics 36, 311–316.

123. Abdennur, N., Goloborodko, A., Imakaev, M., Kerpedjiev, P., Fudenberg, G., Oullette, S., Lee, S., Strobelt, H., Gehlenborg, N., and Mirny, L.A. (2025). cooler (Zenodo) 10.5281/ZENODO.16294112.

124. Imakaev, M., Fudenberg, G., McCord, R.P., Naumova, N., Goloborodko, A., Lajoie, B.R., Dekker, J., and Mirny, L.A. (2012). Iterative correction of Hi-C data reveals hallmarks of chromosome organization. Nat. Methods 9, 999–1003.

125. Open2C, Abdennur, N., Abraham, S., Fudenberg, G., Flyamer, I.M., Galitsyna, A.A., Goloborodko, A., Imakaev, M., Oksuz, B.A., Venev, S.V., et al. (2024). Cooltools: Enabling high-resolution Hi-C analysis in Python. PLoS Comput. Biol. 20, e1012067.

126. Chang, L.-H., Ghosh, S., and Noordermeer, D. (2020). TADs and their borders: Free movement or building a wall? J. Mol. Biol. 432, 643–652.

127. Harris, C.R., Millman, K.J., van der Walt, S.J., Gommers, R., Virtanen, P., Cournapeau, D., Wieser, E., Taylor, J., Berg, S., Smith, N.J., et al. (2020). Array programming with NumPy. Nature 585, 357–362.

128. The pandas development team (2023). pandas-dev/pandas: Pandas (Zenodo) 10.5281/ZENODO.7549438.

129. Virtanen, P., Gommers, R., Oliphant, T.E., Haberland, M., Reddy, T., Cournapeau, D., Burovski, E., Peterson, P., Weckesser, W., Bright, J., et al. (2020). SciPy 1.0: fundamental algorithms for scientific computing in Python. Nat. Methods 17, 261–272.

130. Open2C, Abdennur, N., Fudenberg, G., Flyamer, I.M., Galitsyna, A.A., Goloborodko, A., Imakaev, M., and Venev, S. (2024). Bioframe: operations on genomic intervals in Pandas dataframes. Bioinformatics 40. 10.1093/bioinformatics/btae088.

131. Huey, J.D., and Abdennur, N. (2024). Bigtools: a high-performance BigWig and BigBed library in Rust. Bioinformatics 40. 10.1093/bioinformatics/btae350.

132. Langer, C.C.H., Mitter, M., Stocsits, R.R., and Gerlich, D.W. (2023). HiCognition: a visual exploration and hypothesis testing tool for 3D genomics. Genome Biol. 24, 1–19.

133. Thorvaldsdóttir, H., Robinson, J.T., and Mesirov, J.P. (2013). Integrative Genomics Viewer (IGV): high-performance genomics data visualization and exploration. Brief. Bioinform. 14, 178–192.

134. Robinson, J.T., Thorvaldsdóttir, H., Winckler, W., Guttman, M., Lander, E.S., Getz, G., and Mesirov, J.P. (2011). Integrative genomics viewer. Nat. Biotechnol. 29, 24–26.

135. Kerpedjiev, P., Abdennur, N., Lekschas, F., McCallum, C., Dinkla, K., Strobelt, H., Luber, J.M., Ouellette, S.B., Azhir, A., Kumar, N., et al. (2018). HiGlass: web-based visual exploration and analysis of genome interaction maps. Genome Biol. 19, 125.

136. Hunter, J.D. (May-June 2007). Matplotlib: A 2D Graphics Environment. Comput. Sci. Eng. 9, 90–95.

137. Waskom, M. (2021). seaborn: statistical data visualization. J. Open Source Softw. 6, 3021.

138. jupyterlab: JupyterLab computational environment (Github).

